# Genetic and metabolic regulation of *Mycobacterium tuberculosis* acid growth arrest

**DOI:** 10.1101/186551

**Authors:** Jacob J. Baker, Robert B. Abramovitch

## Abstract

*Mycobacterium tuberculosis* (Mtb) senses and adapts to acidic environments during the course of infection. Acidic pH-dependent adaptations include the induction of metabolic genes associated with anaplerosis and growth arrest on specific carbon sources. In this study, reverse and forward genetic studies were undertaken to define new mechanisms underlying pH-dependent adaptations. Here we report that deletion of isocitrate lyase (*icl1/2*) or phosphoenolpyruvate carboxykinase (*pckA*) results in reduced growth at acidic pH and altered metabolite profiles, supporting that remodeling of anaplerotic metabolism is required for pH-dependent adaptation. Mtb cultured at pH 5.7 in minimal medium containing glycerol as a single carbon source exhibits an acid growth arrest phenotype, where the bacterium is non-replicating but viable and metabolically active. The bacterium uptakes and metabolizes glycerol and maintains ATP pools during acid growth arrest and becomes tolerant to detergent stress and the antibiotics isoniazid and rifampin. A forward genetic screen identified mutants that do not arrest their growth at acidic pH, including four enhanced acid growth (*eag*) mutants with three distinct mutations in the PPE gene MT3221. Overexpression of the MT3221(S211R) variant protein in wild type Mtb results in enhanced acid growth and reduced drug tolerance. Together, these findings provide new evidence for a genetic and physiological basis for acid growth arrest and support that growth arrest is an adaptive process and not simply a physiological limitation associated with acidic pH.

**Author Summary:** The bacterium *Mycobacterium tuberculosis* (Mtb) causes the disease tuberculosis in humans. During infection Mtb colonizes a variety of environments that have acidic environments and Mtb must adapt to these environments to cause disease. One of these adaptations is that Mtb slows and arrests its growth at acidic pH, and the goal of this study was to examine the genetics and physiology of these pH-dependent adaptations. We found that Mtb modifies its metabolism at acidic pH and that these adaptations are required for optimal growth. We also found that acidic pH and specific nutrient sources can promote the bacterium to enter a state of dormancy, called acid growth arrest, where the bacterium becomes tolerant to antibiotics. Mutants were identified that do not arrest their growth at acidic, revealing that acid growth arrest is a genetically controlled process. Overall, understanding how Mtb adapts to acidic pH has revealed pathway that are required for virulence and drug tolerance and thus may identify new targets for drug development that may function to shorten the course of TB therapy.

## Introduction

The success of *Mycobacterium tuberculosis* (Mtb) as a human pathogen can be attributed in part to its ability to slow and arrest its growth in response to immune pressures. Upon infection of humans, Mtb survives long periods of slowed and arrested growth, remaining quiescent for decades before reactivating to cause disease [1]. Even in active cases of tuberculosis, the long course of antibiotic treatment required to clear infection, 6 to 9 months [2], is understood to be necessary due to the phenotypic tolerance of Mtb subpopulations with reduced growth [3]. The issue of phenotypic tolerance is exacerbated by the diversity of Mtb-containing lesions that develop during infection, implying that throughout infection Mtb populations exist in a variety of host environments permissive for varying degrees of growth [4]. Even from sputum samples of infected patients, Mtb can be isolated that displays increased tolerance to front line Mtb antibiotics [5]. Thus, efforts to understand how Mtb regulates growth during infection are relevant to the successful treatment of tuberculosis.

While the exact mechanisms responsible for growth arrest *in vivo* are not completely understood, several studies have investigated the response of Mtb to host-relevant stresses *in vitro.* In response to environmental conditions such as hypoxia or starvation, Mtb enters a state of non-replicating persistence (NRP) [6,7]. These *in vitro* persistent states are characterized by metabolic remodeling and increased antibiotic tolerance [8-10]. Similar observations of antibiotic tolerance have been made during Mtb growth arrest in response to nitric oxide [11], low iron [12], and in multiple stress models [13]. The relationship between non-replicating persistence and Mtb phenotypic tolerance makes inhibition of Mtb persistence an inviting therapeutic target [14,15].

In addition to these models of Mtb growth arrest, we have previously studied the *in vitro* response of Mtb to the stress of acidic pH. One of the earliest cues encountered by the bacterium during infection is the acidic pH of the macrophage phagosome [16]. Mtb is capable of resisting acid stress, and can maintain viability in cultures as acidic as pH 4.5 [17,18]. This ability to resist acid stress was shown to be impaired in a transposon mutant containing an insertion in Rv3671c *(marP)*, encoding a membrane-associated protein [17]. Notably, this mutant was severely attenuated during murine infection [17]. Additionally, using a zebrafish-*Mycobacterium marinum* infection model, *marP* was shown to be specifically required to survive within the phagolysosome of the host [19], further supporting the hypothesis that the ability of Mtb to resist acid stress is required during infection. In addition to acid resistance, Mtb also responds transcriptionally to the acidic pH of the macrophage within 20 minutes of phagocytosis [16], and deletion of the acid-induced *phoPR* two component regulatory system leads to attenuation [20-22], emphasizing that how Mtb adapts to acidic pH is relevant to its pathogenesis.

We have previously identified that in response to acidic pH, Mtb exhibits carbon source specific growth arrest [23]. Using minimal medium supplemented with single carbon sources, we observed that Mtb arrests growth at pH 5.7 when supplied with most carbon sources. However, the carbon sources cholesterol, acetate, pyruvate, and oxaloacetate permit growth at pH 5.7. The carbon sources that permit Mtb growth at acidic pH feed the anaplerotic node of metabolism. The anaplerotic node is thought to be a key metabolic switch point in the regulation of anabolism, catabolism, and energy production, making proper function of this node important during metabolic adaptation [24]. Transcriptional profiling of Mtb at acidic pH shows the induction of several genes involved in metabolism at the anaplerotic node, including those coding for phosphoenolpyruvate carboxykinase *(pckA)*, isocitrate lyase *(icl1/2)*, malic enzyme *(mez)*, and pyruvate phosphate dikinase *(ppdk)* [23], suggesting that Mtb increases metabolism via the anaplerotic node at acidic pH. Other studies have similarly observed that anaplerotic metabolism is associated with conditions of slowed growth, such as hypoxia [10,25], controlled growth in a chemostat [26], and growth in THP-1 cells [27]. We hypothesized that proper function of the anaplerotic node is required for metabolic adaptation to acid pH. Consistent with this hypothesis, the isocitrate lyase *(icl)* inhibitor 3-NP reduces Mtb growth at acidic pH with pyruvate as the carbon source [23], suggesting that *icl* may be a component of anaplerotic adaptation at acidic pH. Notably, mutants in *icl* and *pckA* are attenuated during animal infection [28-30], supporting that these pH-dependent metabolic adaptations may contribute to Mtb pathogenesis.

The goal of this study is to further define the genetic and physiological basis of acid growth arrest. Given that only carbon sources that feed the anaplerotic node promote Mtb growth and that genes of the anaplerotic node are induced transcriptionally at acidic pH, we investigated and identified a role for *icl/2* and *pckA* in the regulation of Mtb growth and metabolism at acidic pH. Additionally, the discovery of arrested growth at acidic pH on specific carbon sources enabled an unbiased forward genetic screen to define new pathways controlling acid growth arrest. This screen identified that the PPE gene MT3221 is required to arrest Mtb growth at acidic pH. Findings from this study support that acid growth arrest is a genetically controlled physiology that enables the bacterium to tolerate environmental stresses and antibiotics.

## Results

### Role of anaplerotic metabolism in Mtb growth at acidic pH

Given the induction of anaplerotic genes *icl* and *pckA* at acidic pH, as well as the ability of carbon sources of the anaplerotic node to promote growth at acidic pH, we hypothesized that genes regulating anaplerotic metabolism play a role in controlling Mtb growth at acidic pH. To test this hypothesis, we examined the growth of two Mtb mutants with deletions of either *pckA* or *icl1/2*. *ΔpckA* and *Δicl1/2* grew similarly to wild type (WT) Mtb in the rich medium buffered to pH 7.0 (Figure 1). However, growth of both mutants was reduced compared to WT when the rich medium was buffered to pH 5.7, supporting the hypothesis that both *pckA* and *icl1/2* play a role in pH-dependent growth adaptations. To better understand the underlying nature of the reduced growth observed at acidic pH, growth curves were also performed in defined minimal medium buffered at either pH 7.0 or pH 5.7. The *ΔpckA* mutant had reduced growth compared to WT Mtb on glycerol at pH 7.0 and maintained growth arrest at pH 5.7 (Figure S1). The mutant was also unable to grow on pyruvate, acetate, or succinate at either neutral or acidic pH (Figure S1), with OD_6_00 decreasing over the course of the experiment. This growth defect of the *ΔpckA* mutant on non-glycolytic carbon sources is consistent with what has been previously observed in unbuffered media in the H37Rv strain background, and has been linked to a defect in gluconeogenesis [30]. To circumvent this limitation of the *ΔpckA* mutant, growth curves were repeated with the addition of glycerol to the culture media. Addition of glycerol was sufficient to restore growth of the *ΔpckA* mutant on pyruvate and acetate at both pH 7.0 and pH 5.7 (Figure S1). Additionally, culture of the *ΔpckA* mutant in the presence of both glycerol and succinate restored the WT phenotype: growth at pH 7.0 and growth arrest at pH 5.7 (Figure S1). These results suggest that, in addition to the growth defect previously observed in unbuffered medium [30], *pckA* is also required for growth on non-gluconeogenic carbon sources at acidic pH.

**Figure 1.**
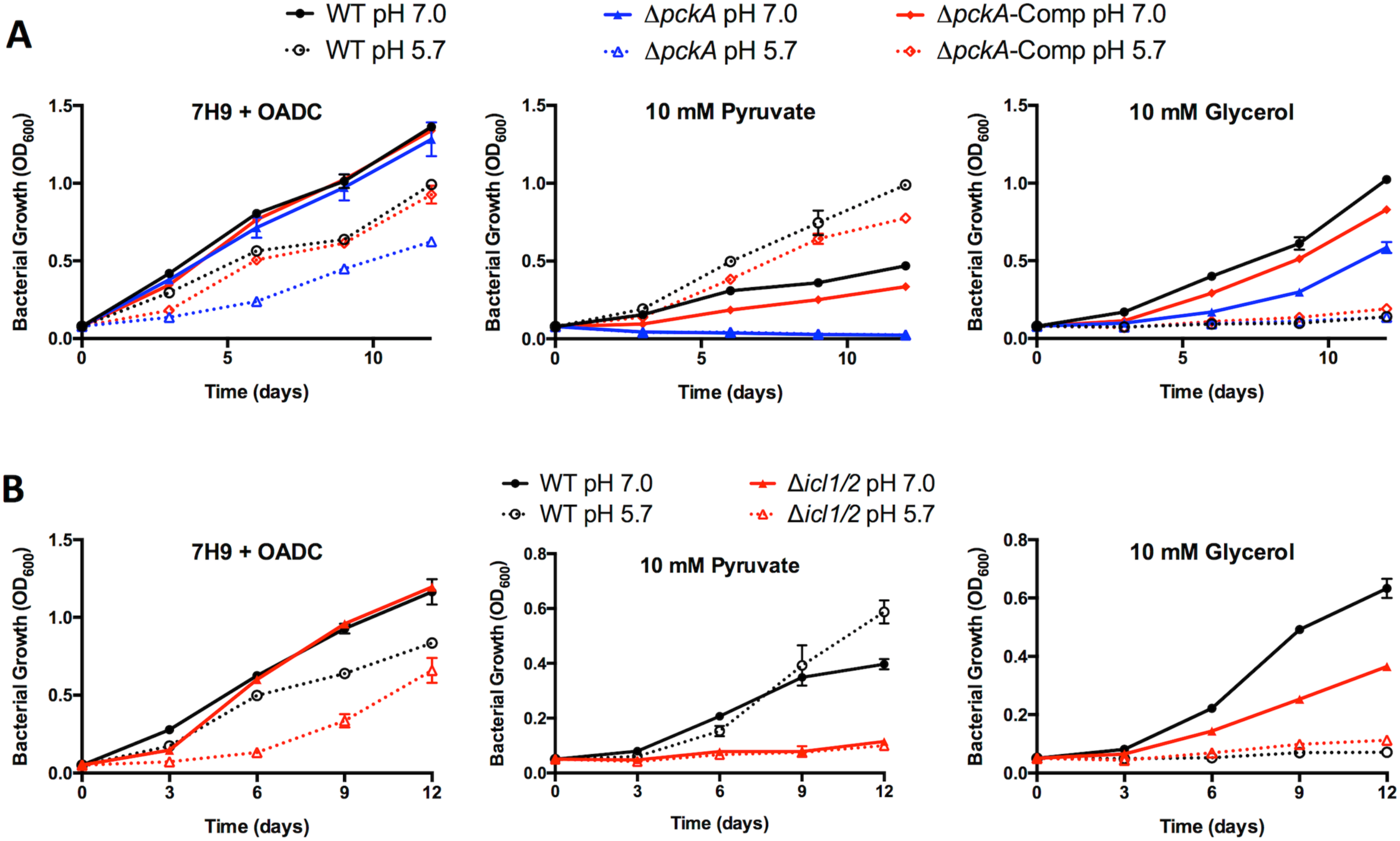
*ΔpckA* and *Δicl1/2* mutants exhibit altered growth profiles at acidic pH and in minimal media. **A.** Growth of CDC1551 WT, *ΔpckA*, and *ΔpckA* complemented (A*pc*fc*A*-Comp) strains in the rich medium 7H9 + OADC and in minimal medium containing either pyruvate or glycerol as a single carbon source, buffered to pH 7.0 or pH 5.7. The *ΔpckA* strain exhibits reduced growth at pH 5.7 in rich medium compared to the WT and complemented strains, and in minimal medium supplemented with pyruvate the OD of *ΔpckA* cultures decreases over time, consistent with bacterial lysis. *ΔpckA* growth on glycerol is reduced compared to the WT and complemented strains at pH 7.0, and is arrested for growth like the WT and complemented strains at pH 5.7. **B.** Growth of Erdman WT and *Δicl1/2* mutant strains in rich and minimal media buffered to pH 7.0 and pH 5.7. The *Δicl1/2* strain exhibits reduced growth at pH 5.7 in rich medium compared to the WT. In minimal medium supplemented with pyruvate, growth is reduced at both pH 7.0 and pH 5.7 in the *Δicl1/2* mutant.

Growth curves in defined minimal media were also performed with the *Δicl1/2* mutant. The *Δicl1/2* mutant exhibited slowed growth on pyruvate at both neutral and acidic pH compared to WT Mtb (Figures S2), only increasing OD_600_ ∼2-fold over the 12-day growth curve, suggesting that anaplerosis via isocitrate lyase is necessary for optimal growth in minimal medium on pyruvate. Growth of the *Δicl1/2* mutant was absent with acetate as the single carbon source (Figure S2), as has been observed previously given the requirement of isocitrate lyase for acetate assimilation [31]. Supplementation of glycerol did not restore WT growth phenotypes in the *Δicl1/2* mutant cultured in minimal medium with pyruvate or acetate (Figure S2), suggesting that the slowed and absent growth observed in the *Δicl1/2* mutant is not due to a deficiency in gluconeogenesis. Although reduced for growth with glycerol as a single carbon source at pH 7.0, at pH 5.7 the *Δicl1/2* mutant exhibited a ∼2-fold increase in OD_600_ beginning after day 3 (Figures 2A, S2A), which is similar to the level of growth observed with pyruvate as a single carbon source. This small amount of growth in conditions typically restrictive for Mtb growth was significantly different from that observed for WT Mtb and was observed in four separate experiments, and we hypothesize that this growth may represent an inability of the *Δicl1/2* mutant to maintain growth arrest at acidic pH. Together, these results indicate that isocitrate lyase is required for optimal growth in minimal medium on pyruvate, and at acidic pH achieves a comparable level of slow growth on both pyruvate and glycerol as single carbon sources.

**Figure 2.**
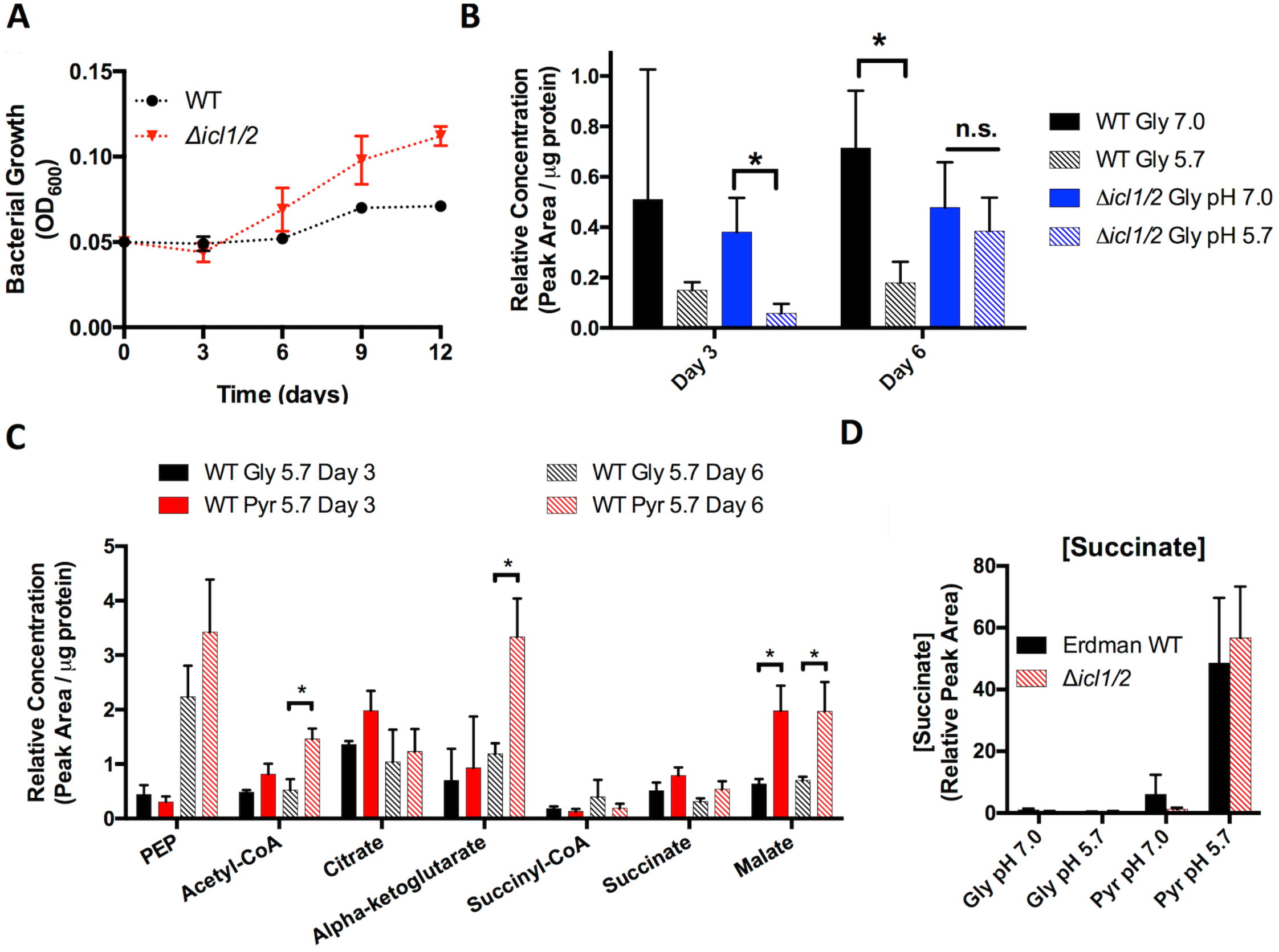
Changes in metabolic profile associated with growth at acidic pH. A. The *Δicl1/2* mutant incubated in glycerol at pH 5.7 exhibits arrested growth at day 3 but enhanced growth at day 6 as compared to the WT. **B.** Relative concentration of succinyl-CoA in WT and *Δicl1/2* mutant in minimal medium with glycerol as a single carbon source buffered to pH 7.0 or pH 5.7. Acidic pH is associated with reduced succinyl-CoA levels in WT Mtb. The *Δicl1/2* mutant exhibited decreased succinyl-CoA levels at day 3, but by day 6 there was no difference in metabolite concentration between pH 7.0 and 5.7. **C.** Relative concentration of selected central carbon metabolites at acidic pH with either glycerol (Gly) or pyruvate (Pyr) as the single carbon source. Mtb has increased concentrations of acetyl-CoA and a-ketoglutarate at day 6, and increased concentrations of malate on day 3 and day 6. **D.** Succinate concentration in supernatant after 3 days culture in minimal medium with Gly or Pyr as single carbon sources buffered to pH 7.0 or pH 5.7. Mtb secretes succinate specifically at pH 5.7 with pyruvate as a single carbon source. Similar levels of succinate accumulation in the supernatant are observed in the *Δicl1/2* mutant.

Given the role of *icl1/2* in both the glyoxylate shunt and the methylcitrate cycle, we sought to clarify which of these activities was responsible for the growth phenotypes observed in minimal media. It has been shown previously that the supplementation of vitamin B12 allows for detoxification of propionyl-CoA via the methylmalonyl pathway [32], and that this supplementation is sufficient to restore growth defects caused by the toxic accumulation of methylcitrate cycle intermediates which are caused by a defective methylcitrate cycle [31,32]. Growth of the *Δicl1/2* mutant in minimal media containing glycerol or pyruvate was not changed with supplementation of vitamin B12 (Figure S3), suggesting that the observed growth phenotypes in the *Δicl1/2* mutant are not due to methylcitrate toxicity and thus may be driven by a dependence of the glyoxylate shunt at acidic pH. In summary, deletion of two enzymes of the anaplerotic node, *pckA* and *icl1/2*, revealed that proper function of the anaplerotic node is necessary for optimal growth of Mtb at acidic pH in rich medium. Furthermore, investigating the growth profiles of these mutants in minimal media has illustrated the predicted importance of anaplerosis for growth in minimal media environments, as both mutants exhibited either slowed growth or growth defects at both neutral and acidic pH.

### Mtb exhibits altered central carbon metabolism at acidic pH

Given the observed role of the anaplerotic node in acidic pH growth regulation as well as the carbon source-specific requirements for growth at acidic pH, we hypothesized that Mtb would exhibit metabolic remodeling at acidic pH that requires proper function of enzymes of the anaplerotic node. To test this hypothesis, metabolic profiling of 12 central carbon metabolism metabolites was performed (Figure S4 and S5). WT Mtb Erdman, an Erdman *Δicl1/2* mutant strain, WT Mtb CDC1551, and a CDC1551 *ΔpckA* mutant strain, were grown on filters placed on agar plates containing minimal medium buffered to either pH 7.0 or pH 5.7 and supplemented with either glycerol or pyruvate as a single carbon source. Metabolites were extracted after 3 and 6 days of culture. Due to the apparent toxicity observed in the *ΔpckA* mutant when cultured with pyruvate as a single carbon source (Figure S1), glycerol and pyruvate were supplemented together for this mutant as well as its WT control. Relative metabolite concentrations were quantified and the data were analyzed for statistical significance using MANOVA followed by post hoc pairwise comparisons that were Bonferroni adjusted to correct for false discovery rate (Supplemental Table 1).

### Decreased succinyl-CoA pools as a biomarker for slowed Mtb growth at acidic pH

Mtb cultured at pH 5.7 with glycerol as a single carbon source exhibits decreased pools of the oxidative TCA cycle intermediate succinyl-CoA (Figure 2B). This decrease was present in both CDC1551 and Erdman WT strains (Figure S4 and S5). Interestingly, it was observed that while succinyl-CoA concentration at pH 5.7 in the *Δicl1/2* mutant is reduced at day 3 (when the mutant is growth arrested), by day 6 (when the mutant exhibits low-level growth on glycerol at pH 5.7, Figure 2A) the succinyl-CoA pools were the same as those observed at pH 7.0 (Figure 2B). These observations suggest that decreased succinyl-CoA may represent a biomarker for growth arrest at pH 5.7, and that isocitrate lyase may play a role in maintaining decreased succinyl-CoA levels at acidic pH.

### Succinate secretion during acidic pH growth

The decreased succinyl-CoA pools during acid growth arresting conditions were also observed at acidic pH with pyruvate as the carbon source, a growth permissive condition (Figure S4 and S5). This observation suggests that the growth observed in pyruvate at pH 5.7 does not require maintaining succinyl-CoA levels, and that an alternative mechanism exists for Mtb growth on pyruvate at acidic pH. While metabolic profiling reveals that WT Mtb exhibits increased pools of phosphoenolpyruvate, acetyl-CoA, and **a**-ketoglutarate by day 6 (Figure 2C), each of these changes are absent at day 3 even though Mtb is already growing at this time point. However, it was observed that at both day 3 and day 6, Mtb does accumulate malate at acidic pH (Figure 2C). This suggested that other time-dependent metabolic adaptations may occur at acidic pH. To identify other time-dependent metabolic adaptations at acidic pH metabolite concentrations in culture supernatants were also measured. Interestingly, succinate accumulated >50-fold in the supernatant specifically in Mtb cultured at pH 5.7 with pyruvate as a single carbon source (Figure 2D). The secretion of succinate was still present in the *Δicl1/2* mutant (Figure 2D), suggesting that Mtb does not require the glyoxylate shunt to secrete succinate. This finding reveals that growth on pyruvate at acidic pH is associated with distinct metabolic adaptations.

Mtb also secretes succinate under conditions of hypoxia [8,33], and it has been hypothesized that succinate secretion allows Mtb to maintain membrane potential in the absence of respiration [8]. This hypothesis is supported by the observation that under conditions of hypoxia the addition of nitrate as an alternate electron acceptor decreases succinate secretion and restores ATP levels and viability to near aerobic levels [33]. To test whether nitrate acts as a similar modulator of respiration at acidic pH, succinate secretion and growth of Mtb was measured with or without addition of sodium nitrate. Interestingly, although having no effect on Mtb growth on pyruvate at pH 7.0, addition of nitrate at pH 5.7 decreased Mtb growth ∼50% (Figure S6A). Succinate secretion by Mtb at pH 5.7 in the presence of nitrate was decreased ∼40% (Figure S6B), although this is likely is secondary to the decreased growth rate, as there are fewer bacteria contributing to secreted succinate pools. The ability of nitrate to modulate Mtb growth at acidic pH but not at neutral pH suggests that Mtb at acidic pH exhibits altered respiration that is responsive to changes in available electron acceptors like nitrate. However, the inability of nitrate to modulate succinate secretion, as is observed under conditions of hypoxia, suggests that this altered respiration is unique from that encountered under conditions of hypoxia.

### *>pckA* mediates transient metabolic adaptations at acidic pH

Despite the *ΔpckA* mutant having no change in growth phenotypes compared to WT Mtb in the culture conditions where metabolic profiling was performed, significant differences in metabolite pools were observed specifically at acidic pH. Compared to WT Mtb, the *ΔpckA* mutant had significantly increased intracellular concentrations of the TCA cycle intermediates citrate, succinate, and malate when cultured at pH 5.7 with glycerol or glycerol and pyruvate as the carbon sources (Figure 3A). The increased concentration of these metabolites was most notable at day 3, and returned toward WT levels by day 6 (with the exception of malate). In Mtb cultured with both glycerol and pyruvate, a 4-fold increase in α-ketoglutarate was also observed (Figure 3B). Notably, no increase in citrate, succinate, or α-ketoglutarate was observed in the *ΔpckA* mutant compared to WT Mtb at neutral pH, and the increase in malate pools at pH 7.0 was less pronounced than at pH 5.7 (Figure S5). The increase in succinate, malate, and citrate specifically at acidic pH in the *ΔpckA* mutant demonstrates that *pckA* reduces accumulation of these TCA cycle intermediates at acidic pH. Indeed, the *ΔpckA* mutant has a 2-fold increase in succinate secretion compared to WT Mtb (Figure 3C), perhaps secondary to this inability to reroute TCA cycle intermediates via the gluconeogenic reaction of *pckA*. However, the return of these TCA intermediates toward WT levels by day 6 suggests that Mtb can compensate metabolically for this pH-dependent overaccumulation. Together, these results support the hypothesis that at acidic pH Mtb utilizes *pckA* for metabolic adaptations.

**Figure 3.**
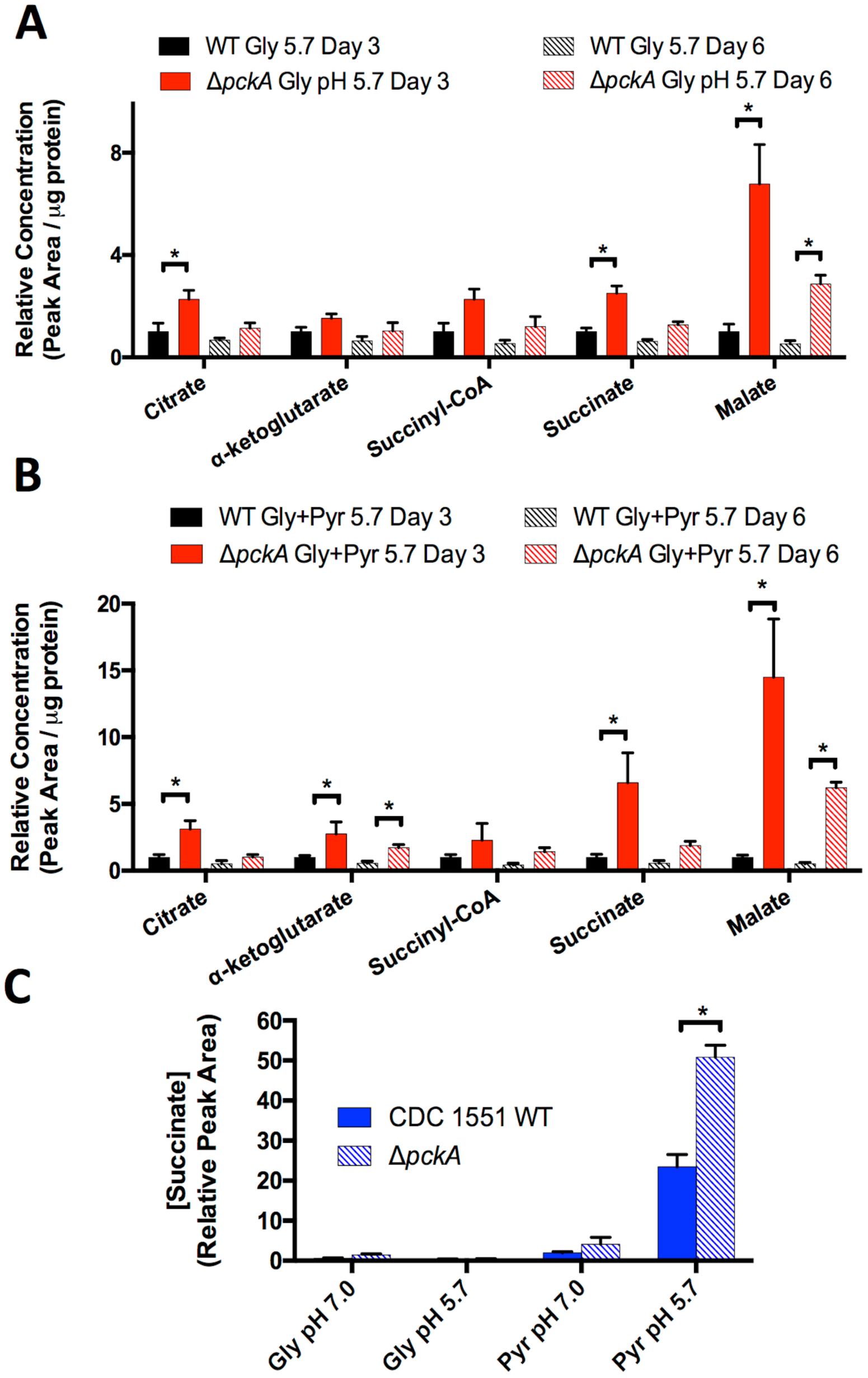
Metabolic profiling of the *ΔpckA* mutant reveals a transient role *for pckA* in metabolic adaptations at acidic pH. Concentration of TCA cycle intermediates in the *ΔpckA* mutant relative to WT Mtb at pH 5.7 with either glycerol (**A**) or glycerol and pyruvate (**B**) as the carbon source(s). Significant differences in the mutant compared to the WT were observed in accumulation of the metabolites citrate, succinate and malate in both carbon sources at day 3, however, the differences resolved by day 3 for citrate and succinate. This finding suggests that initial adaptations to acidic pH may rely upon gluconeogenesis. **C.** Relative concentration of succinate secreted by WT and *ΔpckA* Mtb. The *ΔpckA* mutant exhibits enhanced succinate secretion at Day 3.

Notably, in addition to the increased concentration of citrate at pH 5.7, the *ΔpckA* mutant does not have diminished succinyl-CoA pools at pH 5.7 compared to pH 7.0 at day 3 (Figure S5). By day 6, succinyl-CoA pools in the *ΔpckA* mutant declined to levels similar to the WT (Figure S5). This observation further supports the view that loss of *pckA* leads to a transient disruption of WT metabolic remodeling at acidic pH. However, both the return of metabolite levels toward WT concentrations in the *ΔpckA* mutant by day 6 and the shared growth phenotypes of the mutant and WT Mtb suggest that Mtb can compensate for deletion of *pckA* during metabolic adaptation at acidic pH.

### Growth arrest at acidic pH induces a metabolically active, non-replicating state in Mtb

Previously, we have observed that Mtb cultured at pH 5.7 in minimal medium containing glycerol as a single carbon source is arrested for growth but maintains viability [23]. A long-term viability assay was performed to examine if Mtb remains in a growth arrested state over the course of several weeks. It was observed that Mtb maintains viability in the absence of replication at pH 5.7 for the 39-day duration of the experiment (Figure 4A). As another measure of viability, ATP concentration was monitored. ATP in Mtb cultured at pH 5.7 in glycerol was significantly diminished as compared to pH 7.0 after 6 days of culture, albeit still ∼5-fold higher than that of Mtb at either pH 7.0 or 5.7 after 26 days of culture (Figure 4B), a time point that is assumed to be carbon limited relative to day 6. Together, these results demonstrate that Mtb cultured in minimal medium with glycerol as a single carbon source buffered to pH 5.7 is a viable, non-replicating state, a physiology that will hereafter be referred to as acid growth arrest.

**Figure 4.**
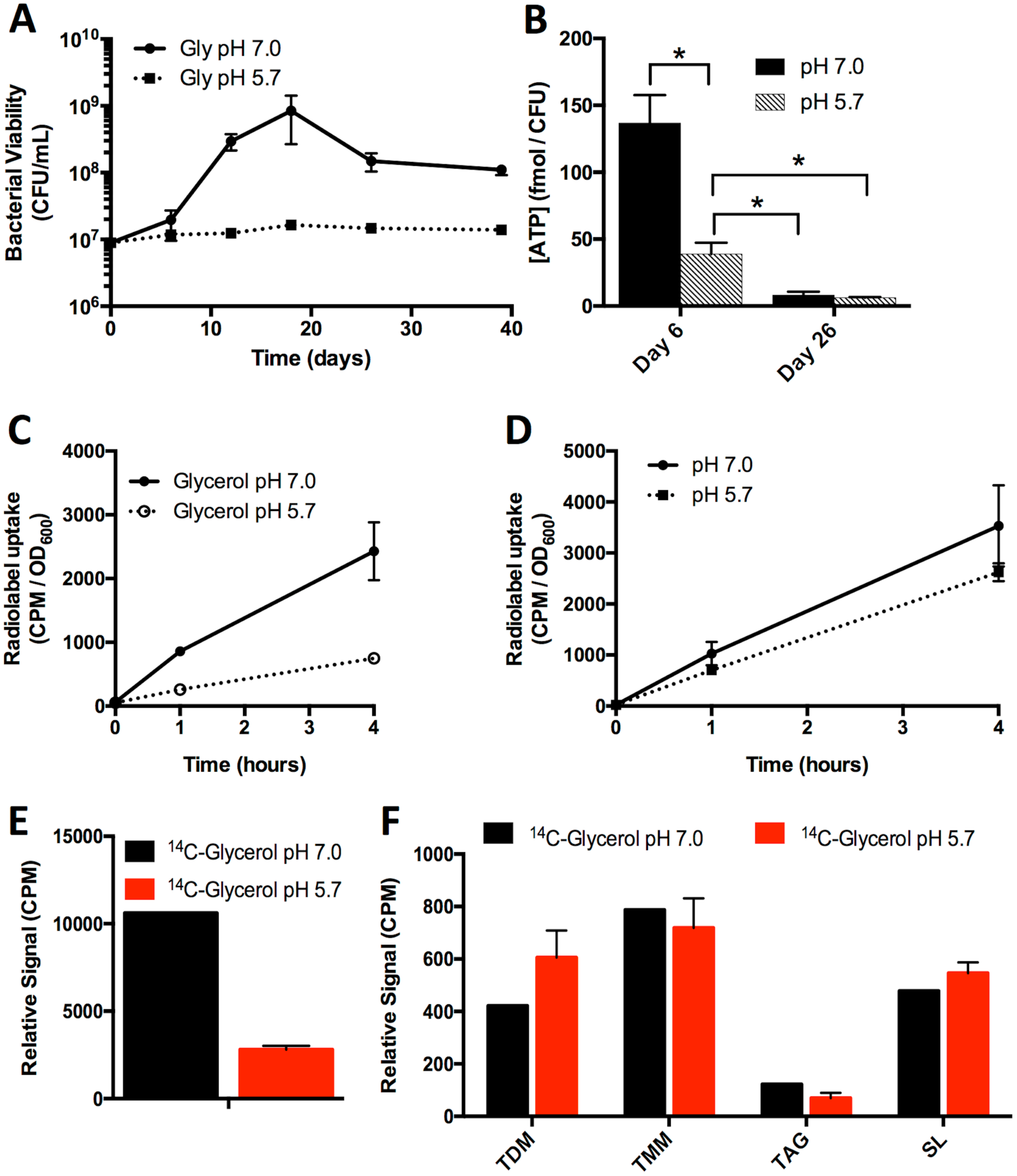
Mtb remains viable and metabolically active during acid growth arrest. **A.** Mtb remains viable during culture in minimal medium buffered to pH 5.7 with glycerol as a single carbon source in the absence of growth. **B.** The concentration of ATP in Mtb under acid growth arrest is reduced compared to pH 7.0, but still higher than that observed on Day 26 at pH 7.0 or pH 5.7. Error bars represent the standard deviation. *p < 0.05, using a Student’s t-test. **C.** Mtb was incubated in minimal medium with glycerol as a single carbon source at pH 7.0 and pH 5.7 and the uptake of ^14^C-glycerol was monitored over time. Mtb uptakes ^14^C-glycerol during acid growth arrest, albeit at a reduced rate. **D.** Uptake of ^14^C-glycerol. Mtb cultures were conditioned to minimal medium with glycerol as a single carbon source for 3 days, pelleted, and resuspended in PBS + 0.05% Tween-80. Accumulation of ^14^C-glyerol was measured over time, with no significant difference in accumulation observed based on two-way ANOVA (p=0.239). Error bars represent standard deviation.**E.** Incorporation of ^14^C-glycerol into Mtb lipids. Following 10 days of culture with ^14^C-glycerol, lipids were extracted and total radioactivity of the samples was measured. Mtb cultured at pH 5.7 had reduced incorporation of ^14^C in lipids, although this difference may be in part attributable to increased bacterial numbers over time at pH 7.0. **F.** Relative radiolabeled lipid species abundance. Thin layer chromatography (TLC) was performed by spotting 5,000 CPM of ^14^C-labelled lipids at the origin and developing the TLC in the necessary solvents for separation of trehalose di‐ and monomycolate (TDM, TMM), and sulfolipid (SL) as described in the methods. For each lipid species, bars indicate relative signal of each lipid species.

To determine whether Mtb under acid growth arrest was still metabolically active, Mtb uptake of ^14^C-glycerol was measured. Over time, Mtb accumulated ^14^C-glycerol at pH 5.7, albeit ∼70% lower than at pH 7.0 (Figure 4C). To account for potential metabolite carryover from rich medium, in a separate experiment Mtb was adapted for 3 days to minimal medium with glycerol as a single carbon source at either pH 7.0 or pH 5.7 and washed immediately prior to the addition of radiolabeled carbon sources. In this case, no significant difference in radiolabel uptake was observed (Figure 4D), suggesting that acid growth arrest is not simply due to changes in cell envelope permeability and glycerol uptake. To determine whether the imported ^14^C-glycerol is metabolized by Mtb under acid growth arrest, the incorporation of ^14^C-glycerol into Mtb lipids was measured. While the level of lipid labeling observed in acid growth arrested Mtb was again reduced ∼70% compared to pH 7.0 (Figure 4E), radiolabel incorporation into trehalose di‐ and monomycolate (TDM, TMM), triacylglycerol (TAG), and sulfolipid (SL) was observed in growth arrested Mtb (Figure 4F, S7). The uptake of glycerol as well as its anabolic incorporation into Mtb lipids suggests that Mtb under acid growth arrest is metabolically active, and supports the view that acid growth arrest is a metabolically active, non-replicating state.

### Acid growth arrest is associated with increased antibiotic and SDS tolerance

A hallmark of metabolically active, non-replicating states is the development of phenotypic tolerance to antibiotics and stress [9,12,34,35]. A role of acidic pH in promoting antibiotic tolerance by preventing cytosolic alkalinization has been previously shown in *Mycobacterium smegmatis* [36]. The sensitivity of Mtb to isoniazid, rifampin, and the detergent sodium dodecyl sulfate (SDS) was measured to test whether acid growth arrest was associated with phenotypic drug or stress tolerance. The concentration of drug necessary to kill 90% of Mtb (MBC_90_) at pH 7.0 and pH 5.7 in both rich and minimal media was determined via spot plating of Mtb treated with different concentrations of drug for 6 days. The MBC_90_ of isoniazid, rifampin, and SDS was increased >5 fold in Mtb under acid growth arrest compared to the other conditions tested, demonstrating that acid growth arrest does increase Mtb tolerance to these drugs (Table 1). Notably, Mtb growing in minimal medium with pyruvate as a single carbon source exhibits a lower MBC_90_ to isoniazid at pH 5.7 compared to pH 7.0; in contrast, tolerance is not observed to rifampin or SDS in pyruvate at pH 5.7. These results indicate that Mtb exhibits phenotypic tolerance during acid growth arrest.

**Table 1.** Minimum bactericidal concentration (MBC_90_) of Isoniazid, Rifampin, and SDS in different culture conditions. MBC_90_ is defined as the concentration of drug necessary to reduce CFU by 90% after 6 days of exposure in the specified culture conditions. The culture conditions tested were rich medium (7H9 + OADC) and minimal medium containing glycerol or pyruvate as a single carbon source, buffered to pH 7.0 or pH 5.7. All culture conditions promote Mtb growth except glycerol as a single carbon source at pH 5.7, which is arrested for growth.

|  | Isoniazid (μM) | Rifampin (μM) | SDS (%) |
| --- | --- | --- | --- |
| Rich Medium pH 7.0 | 3.5 | <0.3 | 0.03 |
| Rich Medium pH 5.7 | 1.6 | <0.3 | 0.03 |
| MMAT Glycerol pH 7.0 | 1.6 | <0.3 | 0.03 |
| **MMAT Glycerol pH 5.7** | **>10** | **1.6** | **0.2** |
| MMAT Pyruvate pH 7.0 | 10 | <0.3 | 0.06 |
| MMAT Pyruvate pH 5.7 | 4 | <0.3 | 0.08 |

### A genetic screen to identify mutants with enhanced acidic pH growth arrest

In other persistent states of Mtb, such as starvation [37] or hypoxia [38], cessation of growth is understood to be due to a physiological limitation, such as absence of a carbon source or a terminal electron acceptor. However, Mtb under acid growth arrest is provided both a metabolically utilized carbon source, glycerol, as well as a terminal electron acceptor, oxygen. We hypothesized that instead of representing a physiological limitation, acid growth arrest is a regulated adaptation of Mtb. To test this hypothesis, a genetic screen was performed to identify mutants unable to arrest growth at acidic pH. A transposon mutant library containing >100,000 mutants was plated on agar plates containing minimal medium buffered to pH 5.7 and supplemented with 10 mM glycerol. Mutants with enhanced acid growth *(eag* mutants) formed colonies that were isolated and confirmed as *eag* mutants by measuring bacterial growth in minimal medium supplemented with 10 mM glycerol at pH 5.7 (Figure S8A). In total, 165 mutants were isolated from plates, of which 98 were confirmed as *eag* mutants. In addition to the transposon (Tn) mutant screen, wild type Mtb was plated on the same acid growth arrest condition to screen for spontaneous *eag* mutants. Two spontaneous mutants were isolated, both exhibiting robust growth in minimal medium supplemented with glycerol at pH 5.7 (Figure 5A).

**Figure 5.**
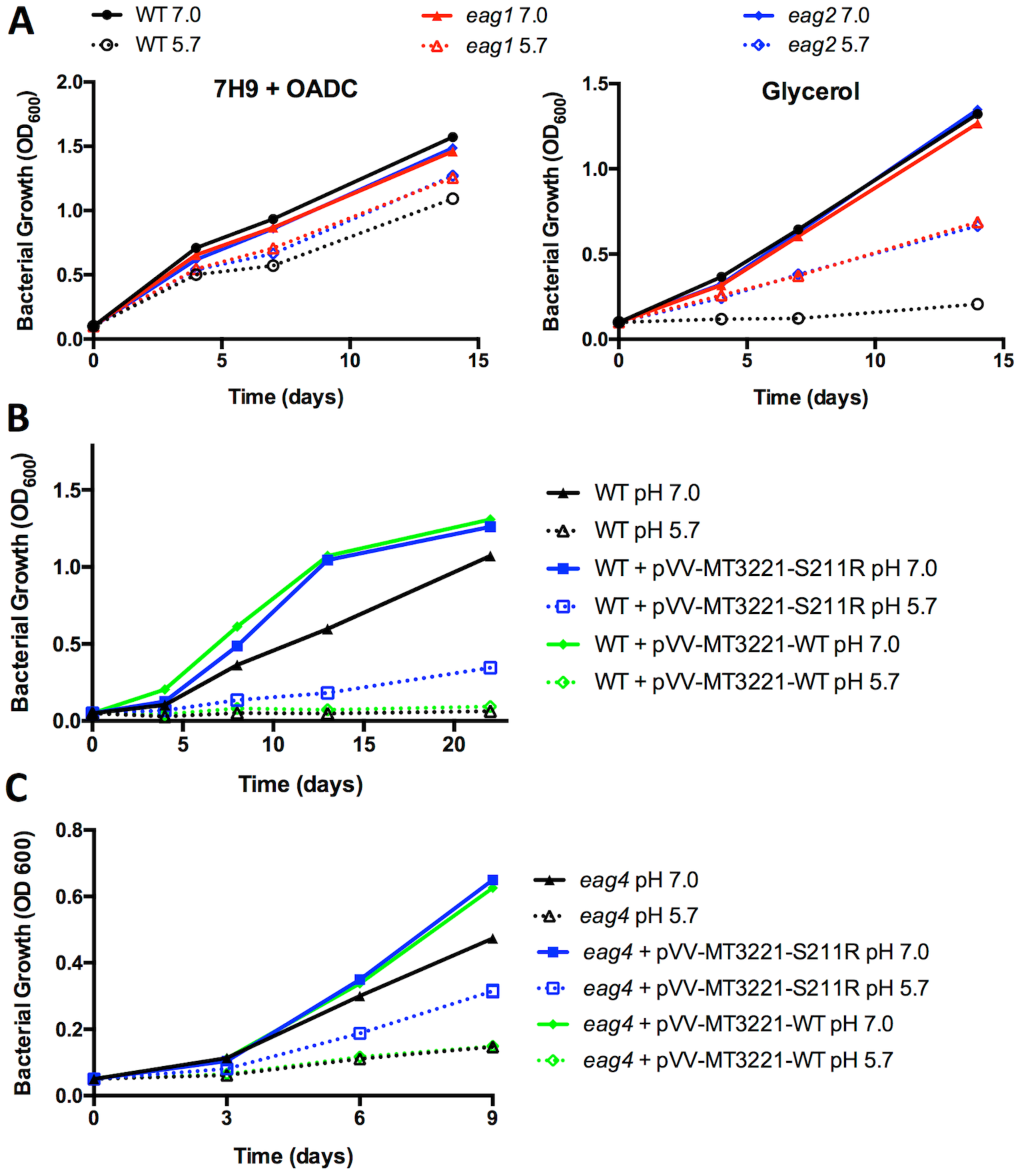
The S211R-encoding mutant allele of MT3221 enhances Mtb growth at acidic pH on glycerol. **A.** Growth curves in rich medium (7H9+OADC) and in minimal medium with glycerol as a single carbon source with two spontaneous *eag* mutants. Notably, the enhanced growth of *eag* mutants is only observed at pH 5.7 and not at pH 7.0. **B.** Growth curve of wild type Mtb (black) and wild type Mtb containing a plasmid overexpressing either the wild type or S211R-encoding mutant allele of MT3221 (blue and green, respectively). Both overexpression constructs increase Mtb growth at pH 7.0, but only the mutant allele promotes Mtb growth at pH 5.7. **C.** Growth of the Mtb *eag4* mutant containing spontaneous S211R-encoding mutation in MT3221. Overexpression of the wild type allele does not arrest growth at pH 5.7, and overexpression of the mutant MT3221 allele increases growth in the *eag4* mutant. Both overexpression constructs increase growth of the *eag4* strain at pH 7.0. Error bars represent the standard deviation.

The Tn insertion site was determined for the confirmed *eag* mutants. For select Tn mutants, complementation was attempted via introduction of an integrative plasmid expressing the wild type version of the disrupted gene with its native promoter. This attempt at complementation did not restore the wild type growth arrest phenotype in the tested mutants. For example, for the *eag4* mutant a Tn insertion was mapped to gene MT1359 and reintroduction of the MT1359 gene and its native promoter on an integrative plasmid did not complement the *eag4* mutant (Figure S8B), even though measurement of transcript levels revealed that the complementation constructs did restore MT1359 mRNA levels of the disrupted genes to wild type levels (Figure S8C). The observation of genetic complementation without phenotypic complementation suggests that Tn disrupted genes were not responsible for the enhanced acid growth phenotype and that the mutant genes driving the observed phenotypes were either the result of spontaneous mutations elsewhere in the genome of polar effects of the Tn insertion.

Given the isolation of spontaneous *eag* mutants, we hypothesized that the lack of complementation in the transposon *eag* mutants may be due to spontaneous mutations within the transposon library. To test this hypothesis, whole genome sequencing was performed on both the spontaneous mutants and select transposon mutants. Of the six *eag* mutants sequenced, four contained a missense mutation in the Mtb gene *PPE51* (MT3221, an orthologue of the H37Rv gene Rv3136), including both spontaneous *eag* mutants (*eag1* and *eag2)* and two Tn mutants *(eag3 and eag4*, Table 2). The presence of single nucleotide variants was confirmed by amplification and Sanger sequencing of the MT3221 gene from each mutant as well as a wild type control. The MT3221 amino acid substitutions include S211G, S211R and A228D showing that the substitutions are clustered in the central region of the 380 amino acid protein. Overexpression of the S211R-encoding mutant allele of MT3221 in wild type Mtb was sufficient to allow growth at acidic pH with glycerol as a single carbon source (Figure 5B), whereas overexpression of the wild type MT3221 allele in the *eag4* mutant background did not restore growth arrest (Figure 5C), revealing that the S211R-encoding mutation has a dominant effect on enhanced growth at acidic pH. The MT3221 overexpression construct was also introduced in the *eag4* mutant. As anticipated, the overexpression of the WT MT3221 allele did not inhibit enhanced acid growth (Figure 5C). Interestingly, overexpression of MT3221-S211R in the *eag4* background led to a more substantial enhanced acid growth phenotype, suggesting that expression levels of the MT3221 could function to regulate growth at acidic pH. Thus, the mutation in MT3221 gene expressing the S211R variant protein was shown to be sufficient for enhanced growth at acidic pH, demonstrating a genetic basis for acid growth arrest.

**Table 2.** Summary of variants identified by whole genome sequencing. Variants were identified using a GATK workflow as described in the experimental methods. 3 unique missense mutations were identified in *MT3221* at 2 unique sites in 4 different mutants. The spontaneous mutants are named *eag1* and *eag2*, whereas the two transposon mutants are named *eag3* and *eag4* with the position of transposon insertion as indicated.

| Strain | Position | Reference | Alternative | Quality score | Substitution | Gene ID | Annotation | Transposon mutant |
| --- | --- | --- | --- | --- | --- | --- | --- | --- |
| *eag1* | 338965 | G | C | 602 | intergenic | Upstream MT0292 | PPE3 |  |
|  | **3497961** | **C** | **A** | **7022** | **A228D** | **MT3221** | **PPE51** |  |
| *eag2* | **3497961** | **C** | **A** | **2462** | **A228D** | **MT3221** | **PPE51** |  |
| *eag3* | **3497909** | **A** | **C** | **7553** | **S211G** | **MT3221** | **PPE51** | *tn::MT1921* at 481 bp |
|  | 3973579 | G | T | 7077 | P131V | MT3646 | Hypothetical protein |  |
| *eag4* | 563721 | C | T | 7272 | R140S | MT0487 | Hypothetical protein | *tn:: MT1359* at 319 bp |
|  | 3242629 | A | C | 74 | H955P | MT3000 | *ppsA* |  |
|  | **3497911** | **C** | **G** | **6310** | **S211R** | **MT3221** | **PPE51** |  |

### eag mutants have reduced phenotypic tolerance

Given the increased tolerance of Mtb during acid growth arrest, we hypothesized that mutants with enhanced acidic pH growth would be more sensitive to both antibiotic and physiological stress. To test this hypothesis, wild type, the *eag4* mutant, and WT Mtb containing a plasmid overexpressing either the wild type or mutant *MT3221* overexpression constructs were treated with isoniazid, rifampin, or PA-824 after 3 days acclimation to minimal medium supplemented with glycerol as a single carbon source buffered to either pH 7.0 or pH 5.7. After exposure to each drug for 6 days, Mtb was plated for viability. Compared to wild type Mtb treated with rifampin, the *eag4* mutant exhibited a 1-log reduction in viability at acidic pH (Figure 6A). Similarly, the *eag4* mutant and wild type Mtb overexpressing the *MT3221*-S211R allele exhibited increased sensitivity to isoniazid at pH 5.7 compared to wild type Mtb, with wild type Mtb overexpressing the wild type MT3221 allele exhibiting sensitivity intermediate to wild type and mutant Mtb (Figure 6B). For both isoniazid and rifampicin, no significant difference in sensitivity was observed between strains at pH 7.0 (Figure 6B). The bicyclic nitroamidazole PA-824 was shown to be comparably effective at killing Mtb at both pH 7.0 and pH 5.7, demonstrating that Mtb growth arrest is not protective against all classes of antibiotics (Figure 6C). The increased sensitivity of mutants with enhanced growth at acidic pH to both rifampin and isoniazid supports the hypothesis that loss of growth arrest at acidic pH increases Mtb antibiotic sensitivity. Furthermore, the lack of tolerance to PA-824 at acidic pH demonstrates the potential of some antibiotics to be effective even during Mtb acid growth arrest.

**Figure 6.**
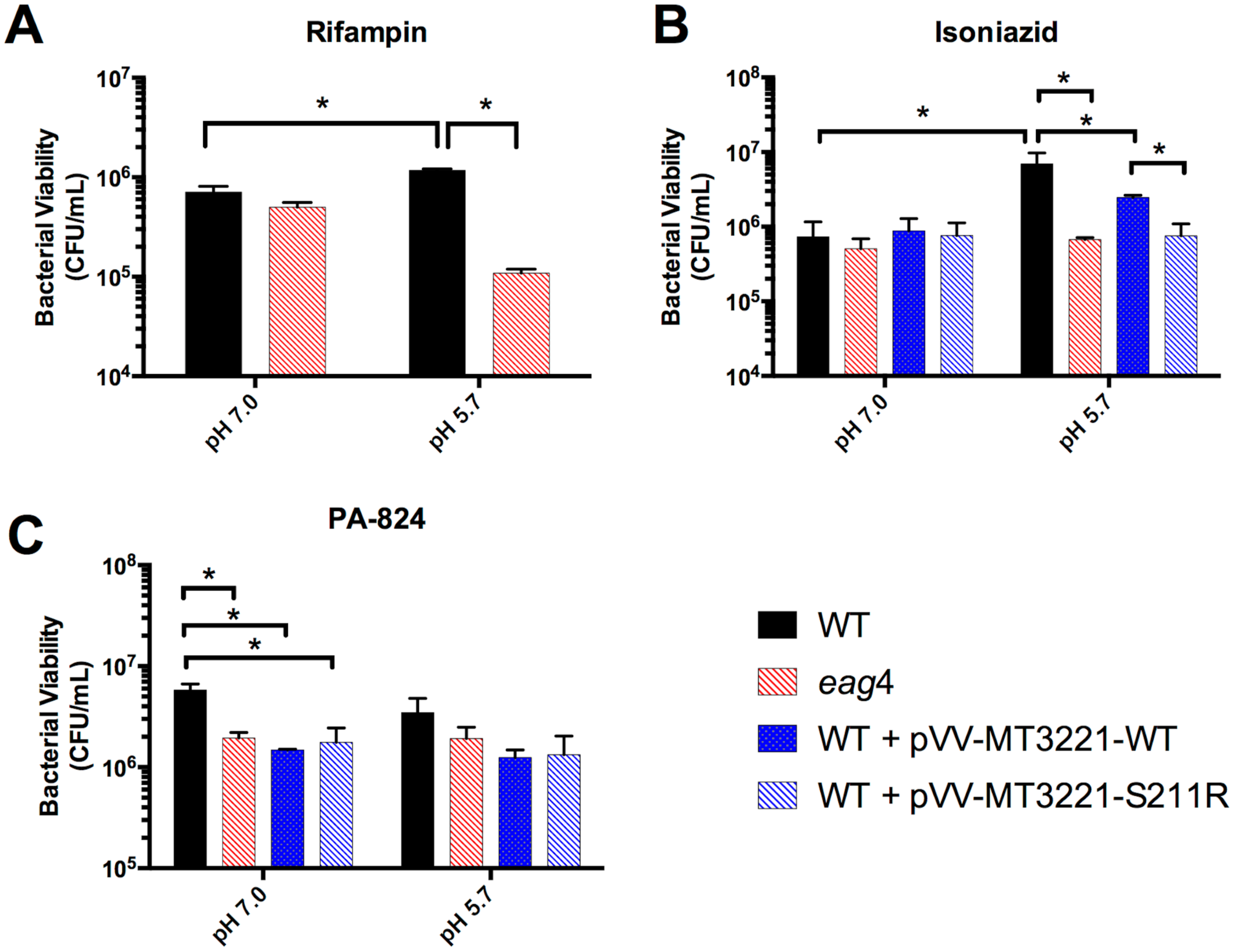
Increased sensitivity of enhanced acid growth mutants to antibiotics. Wild type Mtb (WT), an *eag4* mutant, and WT Mtb expressing either the wild type allele or S211R-encoding mutant allele of *MT3221* (WT + pVV-MT3221-WT and WT + pVV-MT3221-S211R, respectively) were acclimated to minimal medium containing glycerol as a single carbon source for 3 days before adding either rifampin (**A**), isoniazid (**B**), or PA-824 (**C**). The *eag4* mutant exhibited increased sensitivity to both rifampin and isoniazid at pH 5.7 but not pH 7.0, and the strain overexpressing the S211R-encoding *MT3221* mutant allele also had increased sensitivity to isoniazid at pH 5.7. Both the mutant strain and the MT3221 overexpression strains demonstrated increased sensitivity to PA-824 at pH 7.0, but no significant tolerance was observed at pH 5.7. *p < 0.05 based on a student’s t-test.

## Discussion

Mtb exhibits metabolic plasticity in the face of changing environments. Mtb grown under conditions of hypoxia, where redox homeostasis would presumably be difficult to maintain due to the lack of oxygen available for respiration, recycles reduced cofactors via lipid synthesis [12] and increased flux through the reductive TCA cycle [8,33]. With the addition of nitrate, Mtb adapts to another metabolic program during hypoxic culture in which nitrate respiration maintains homeostasis [33,39]. To metabolize branched chain fatty acids and cholesterol, Mtb utilizes *prpCD* and *icl1/2* to prevent toxic accumulation of propionyl-CoA intermediates using the methylcitrate cycle [31,40,41]. The supplementation of vitamin B12 opens yet another pathway for propionyl-CoA metabolism in Mtb, the methylmalonyl pathway [32]. Even under conditions of starvation, Mtb can maintain membrane potential and a basal level of intracellular ATP [9]. Notably, these metabolic adaptations in response to changing environments are crucial to Mtb pathogenesis. Although nonessential for growth *in vitro* in rich medium, several genes encoding for metabolic enzymes have been shown to be required for Mtb survival during infection, including *icl1/2* [28,29], *pckA* [30], *dlaT* [42], and *hoas* [43]. Investigating the mechanisms of metabolic adaptation to environmental stress *in vitro* has helped elucidate the function of these genes during infection as well as explain the basis of their importance in Mtb pathogenesis.

In this study, we sought to define genetic and physiological adaptations to acidic pH, using reverse and forward genetic approaches. Deletion of two genes of the anaplerotic node, *pckA* and *icl1/2*, led to decreased growth at acidic pH but not neutral pH in rich medium, highlighting that Mtb requires anaplerotic metabolism specifically at acidic pH. In minimal media, the main growth phenotypes of both mutants appeared to be related to the inability to perform anaplerosis, as both mutants had difficulty growing in carbon sources other than glycerol, irrespective of pH. The measurement of metabolic intermediates performed here, along with previously published transcriptional profiles [23], provides strong support that Mtb undergoes metabolic remodeling at acidic pH. One of the key outcomes of metabolic remodeling at acidic pH was a decrease in succinyl-CoA levels, which we hypothesize is due to decreased metabolic flux through the oxidative TCA cycle. The oxidative TCA cycle is an energy generating pathway in the form of NADH and ATP [44]. Although other metabolic pathways can similarly generate energy, the oxidative TCA cycle is somewhat unique in that the process requires two steps of irreversible oxidative decarboxylation, from citrate to α-ketoglutarate to succinyl-CoA (Figure 7A). High flux through these irreversible reactions requires that Mtb can incorporate the high-energy cofactors generated into other metabolic reactions. We hypothesize that under stressful conditions where metabolism may be restricted, avoiding flux through the oxidative TCA cycle may prevent Mtb from overaccumulating these intermediates and thus contribute to slowed growth. We observed that in the *Δicl1/2* mutant, increased pools of succinyl-CoA coincided with the loss of growth arrest at acidic pH. This observation supports the hypothesis that *icl1/2* may play an important role in regulating the flux through the oxidative TCA cycle to regulate Mtb growth (Figure 7A). Notably, *icl* induction and the glyoxylate shunt has been associated with other slowed growth conditions including hypoxia [10,25], growth in chemostat [26] and survival in THP-1 cells [27], suggesting that acidic pH-dependent regulation of *icl* and growth may have growth rate controlling interactions when combined with other immune stresses.

**Figure 7.**
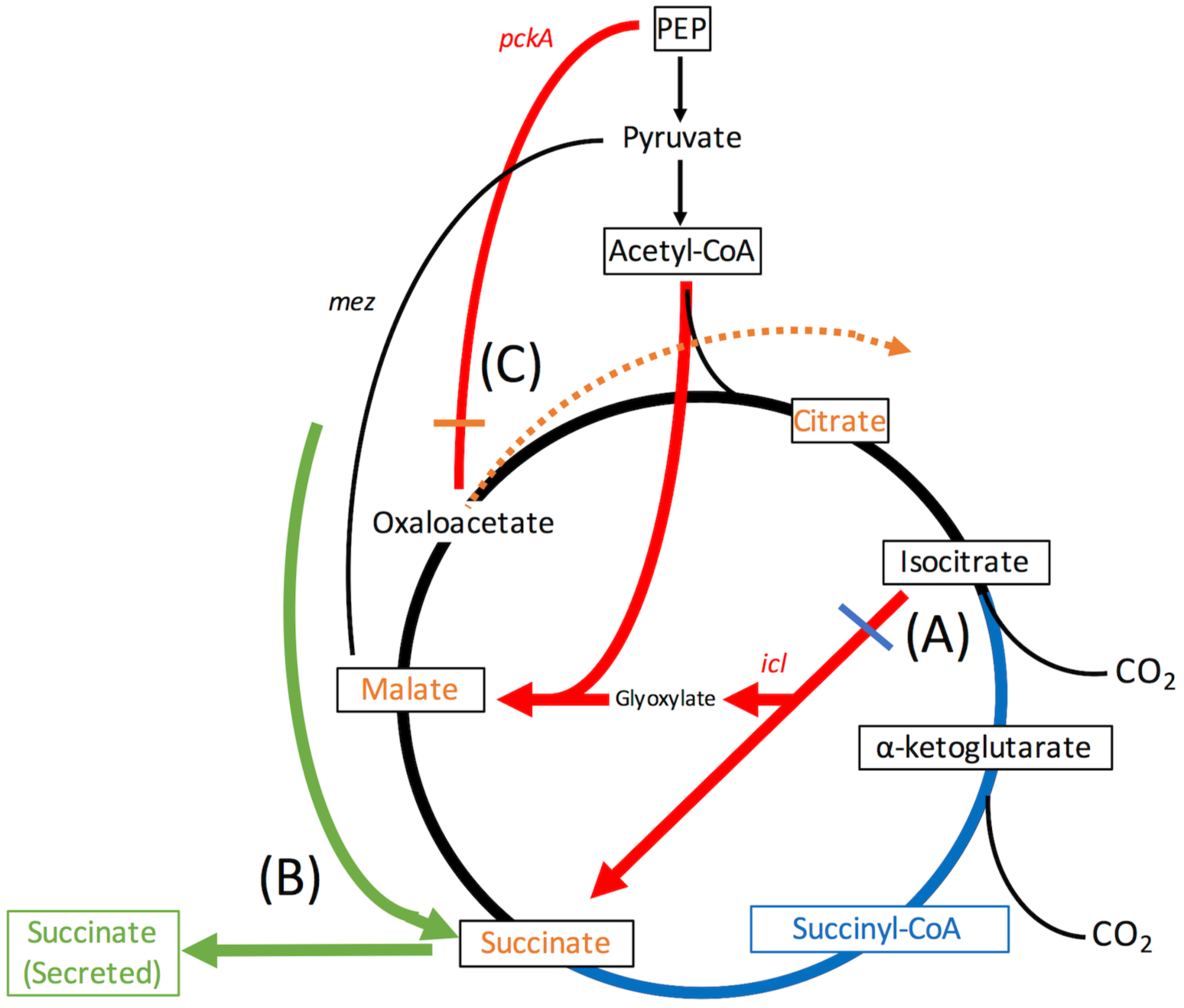
A. Model of metabolic adaptations at acidic pH. **A.** Highlighted in blue, at acidic pH, the transcriptional induction of isocitrate lyase *(icl)* and decreased concentration of succinyl-CoA suggest that Mtb increases metabolic flux through the glyoxylate shunt and decreases flux through the oxidative TCA cycle. This remodeling limits production of NADPH and ATP via irreversible oxidative decarboxylation and additionally increases the production of reductive TCA cycle intermediates (succinate and malate). **B.** Highlighted in green, growth at acidic pH is associated with secretion of succinate. The *Δicl1/2* mutant still secretes succinate, suggesting that Mtb can utilize a route other than the glyoxylate shunt for succinate secretion. **C.** Highlighted in orange, the transcriptional induction of *pckA* at acidic pH, as well as the observed accumulation of succinate, malate, and citrate in the *ΔpckA* mutant at acidic pH suggests that Mtb utilizes the gluconeogenic reaction of *pckA* to divert the increased reductive TCA cycle intermediates away from citrate synthase. The decrease of these intermediates in the *ΔpckA* mutant to WT levels by day 6 suggests that Mtb can compensate metabolically for loss of *pckA*, possibly by succinate secretion.

The identification of succinate secretion in growing Mtb at acidic pH is an unexpected observation (Figure 7B). In conditions of hypoxia, succinate secretion has been shown to be necessary to prevent succinate accumulation secondary to increased utilization of the glyoxylate shunt and reductive TCA cycle for oxidized cofactor regeneration [8,33]. The phenotype in hypoxia has been linked to decreased respiration in hypoxic environments, as the addition of nitrate as an alternative electron acceptor decreased succinate secretion [31]. In Mtb growing at acidic pH, the reason for succinate secretion is less obvious. In this culture condition, oxygen is available to Mtb, suggesting oxidative respiration is possible at acidic pH. However, it was shown previously that at acidic pH Mtb induces transcriptionally type-II NADH dehydrogenase and *bd*-type cytochrome oxidase and represses expression of the type-I NADH dehydrogenase complex and *c*-type cytochrome oxidase [23]. Why Mtb switches from a proton translocating to non-proton translocating electron transport chain is not well understood, but does suggest that changes in respiration may indeed be occurring at acidic pH. Furthermore, while the addition of nitrate as an alternative electron acceptor did not abolish succinate secretion as was observed under hypoxia, it did lead to an acidic pH-specific reduction in growth at acidic pH. Notably, the use of nitrate for respiration at acidic pH has been observed before, with Mtb in a hypoxic and acidic environment requiring nitrate respiration to maintain viability [39]. These observations suggest that changes in Mtb respiration may contribute to the metabolic remodeling observed at acidic pH.

The marked increase in TCA intermediates at acidic pH in the *ΔpckA* mutant suggests an increased requirement for *pckA*-dependent gluconeogenesis at acidic pH (Figure 7C). If Mtb shifts its metabolism away from the oxidative TCA cycle and toward the glyoxylate shunt at acidic pH, this shift would lead to a stoichiometric increase in succinate-fumarate-malate intermediates as no carbon is lost through decarboxylation. In this setting, the gluconeogenic reaction of *pckA* allows for an alternate route of metabolism instead of moving these metabolites back into the oxidative TCA cycle via citrate synthase (Figure 7C). Increased metabolism via *pckA* coupled with the increased use of the glyoxylate shunt, as is suggested by metabolic and transcriptional profiling, is reminiscent of the PEP-glyoxylate cycle previously characterized in *E. coli* grown in limited glucose culture [45]. This PEP-glyoxylate cycle has been proposed to be a means of decoupling catabolism from NADPH production, an outcome that is necessary in situations where little biomass is synthesized [45]. In Mtb at acidic pH, we speculate that one of two scenarios lead to a shift to this PEP-glyoxylate type metabolism. First, Mtb could require this decoupling to adapt to the changes in respiration that occur at acidic pH. Alternatively, in response to acidic pH, Mtb could actively remodel metabolism to slow growth, and the decoupling of catabolism from NADPH production provided by the PEP-glyoxylate cycle could aid in halting biosynthesis.

The association between non-replicative persistence and phenotypic tolerance is well documented in the Mtb literature [9,12,34,46]. Notably, the observed association of *icl* induction and drug tolerance at acidic pH is consistent with the reported role of *icl* dependent metabolic adaptations in promoting drug tolerance [47]. The ability of so many distinct environmental conditions to produce a shared phenotype is particularly intriguing, and suggests that the response of Mtb to these environmental conditions may share common mechanisms of persistence. It is worth noting that specific differences in Mtb antibiotic tolerance do exist between the different *in vitro* persistent states. For example, while both hypoxic and nutrient starved non-replicating Mtb exhibit strong tolerance to isoniazid, the hypoxic non-replicating Mtb is less tolerant than starved Mtb to other antibiotics such as rifampin or streptomycin [9]. Under conditions of acid growth arrest in this study, Mtb exhibits increased tolerance to both isoniazid and rifampin, suggesting shared attributes with non-replicating persistence in response to hypoxia or starvation. Like hypoxia, acidic pH does not confer tolerance to PA-824. Given that PA-824 is thought to poison the electron transport chain by acting as an NO donor [48], the lack of resistance to PA-824 during acid growth arrest supports that under this condition Mtb still requires proper function of the electron transport chain.

In our forward genetic screen for mutants that have enhanced growth at acidic pH, we identified and confirmed over 50 such mutants. The identification of these mutants supports the view that acid growth arrest is a genetically controlled phenotype in Mtb. Using whole genome sequencing, we identified single nucleotide variants in the gene MT3221 (Rv3136 in H37Rv), a gene that encodes the Mtb protein PPE51. Like many PE/PPE proteins, the function of PPE51 is not known, however, previous transcriptional profiling of Mtb by our lab shows that MT3221 is induced at acidic pH in both glycerol and pyruvate and in a *phoP*-dependent manner [23,49]. MT3221 expression has also been shown to be reduced during starvation [37]. The ability of point mutations in the gene MT3221 to increase Mtb growth at acidic pH suggests that these mutations can change the function of PPE51 during acid growth arrest. Given that PPE51 is predicted to be a membrane-associated protein, we speculate that these point mutations may increase Mtb growth by changing the ability of PPE51 to interact with other components of the Mtb cell envelope. Two of the sequenced *eag* mutants did not contain mutations in MT3221, demonstrating that other mechanisms of enhanced acid growth exist at acidic pH. We anticipate that further whole genome sequencing of *eag* mutants will identify additional metabolic and regulatory mechanisms responsible for growth arrest at acidic pH.

Growth arrest in Mtb is often assumed to be a required outcome of the physiological limitations of Mtb. In the absence of oxygen as a terminal electron acceptor, the obligate aerobe Mtb requires non-replicating persistence to maintain redox homeostasis, and the disruption of this process leads to a short period of enhanced growth followed by increased cell death [12]. Similarly, the growth arrest in the Loebel starvation model of persistence is readily attributable to the lack of adequate nutrients for energy generation. In these physiologically limited conditions, the fitness of Mtb is appreciated in its ability to remain viable until the situation improves. In the characterization of acid growth arrest in Mtb, we have demonstrated that growth arrest in this condition is not due to physiological limitation, as we have identified numerous mutants capable of growing where wild type Mtb does not. This suggests that Mtb has evolved to undergo acid growth arrest for reasons beyond specifically surviving the acidic pH environment. We propose that the fitness advantage of Mtb through acid growth arrest is in the ability of acid growth arrest to increase phenotypic tolerance. In this way, Mtb could use the host cue of acidic pH to prepare for the myriad other stresses encountered during infection.

pH-driven adaptation is an attractive target for basic research and new drug development efforts. Indeed, pyrazinamide is most active at acidic pH and is a first line drug to treat TB. Mutants defective in pH-dependent adaptations (*e.g. phoPR*, Rv3671c) are attenuated in mice [17,50] and small molecules targeting these pathways have been discovered [49,51]. Moreover, small molecules inhibiting pH-homeostasis have also been identified [52]. Additionally, it was recently shown that Mtb has enhanced sensitivity to thiol and oxidative stress at acidic pH, indicating that pH-dependent changes in redox homeostasis are vulnerable pathways for new drug development [53]. The discovery that acid growth arrest drives drug tolerance and can be genetically disrupted supports the potential to develop small molecules targeting acid growth arrest.

## Materials and Methods

### Bacterial strains and growth conditions

All Mtb experiments, unless otherwise stated, were performed with Mtb strain CDC1551. The *ΔpckA* mutant was generated by homologous recombination and verified by quantitative real time PCR. Complementation of the mutant was achieved by cloning the *pckA* gene as well as its native promoter (1000 bp upstream) into the integrative plasmid pMV306. The *Δicl1/2* mutant and its WT Erdman control were a generous gift from Prof. John McKinney [28]. Cultures were maintained in 7H9 Middlebrook medium supplemented with 10% OADC and 0.05% Tween-80. All single carbon source experiments were performed in MMAT defined minimal medium as described by Lee *et al.* [54]: 1 g/L KH2PO4, 2.5 g/L Na2PO4, 0.5 g/L (NH4)2SO4, 0.15 g/L asparagine, 10 mg/L MgSO4, 50 mg/L ferric ammonium citrate, 0.1 mg/L ZnSO4, 0.5 mg/L CaCl2, and 0.05% Tyloxapol. Medium was buffered using 100 mM MOPS (pH 6.6-7.0) or MES (pH 5.7-6.5) [55]. For growth experiments, Mtb was seeded in T-25 standing tissue culture flasks in 8 ml of minimal medium at an initial density of 0.05 OD600 and incubated at 37°C and 500 μl samples were removed at each time point for optical density measurements. For viability assays, colony forming units were enumerated on 7H10 + 10% OADC agar plates following plating of serial dilutions in PBS + 0.05% Tween-80.

### Metabolic profiling

For metabolic profiling studies, Mtb cultures were centrifuged, washed with 0.9% saline, and resuspended at a final OD_6_00 of 2. 1 mL of washed Mtb was placed on a membrane filter via vacuum filtration and put on agar plates containing the indicated minimal medium [54]. For extraction, Mtb laden filters were transferred to a 6-well plastic plate, frozen on dry ice, and quenched with 500 μL of methanol chilled on dry ice. 175 μL of water and 25 μL of the internal standard 100 μM succinic acid 2,2,3,3-d_4_ were added to each sample, and bacteria were scraped off the membrane filters with a plastic loop and transferred to a 2 mL tube containing 1.2 mL chloroform. Samples were vortexed at 4°C for 30 minutes, centrifuged for 5 minutes at 15,000 rpm, and the upper aqueous phase was dried under liquid nitrogen and frozen at -80 °C until liquid chromatography/mass spectrometry (LC/MS) analysis. For protein quantification, the interphase of the sample was dried and resuspended by sonication in 20 mM Tris pH 8.0, 0.1 % sodium dodecyl sulfate (SDS), and 6M urea. For LC/MS analysis, samples were resuspended in 10 mM tributylamine (TBA), 10 mM acetic acid (AA), 97:3 water:methanol. Samples were applied to a C18 BEH-amide column and metabolites were separated using a 10 minute inlet method with the mobile phase starting at 99:1 10 mM TBA: 10 mM AA in 97:3 water:methanol and finishing at 100% methanol. Multiple reaction monitoring (MRM) channels were developed for each metabolite using known standards, and metabolites were identified by comparing monoisotopic mass and retention time to these standards.

### Analysis of mycobacterial lipids

For lipid remodeling experiments, bacterial cultures were pre-adapted in MMAT medium with 10 mM glycerol for 3 days, washed and then resuspended in MMAT medium 10 mM glycerol and 8 μCi of [U-^14^C] Glycerol. Following 10 days of labeling, bacteria were pelleted, washed, and the lipids extracted as described previously [23]. Total radioactivity and ^14^C incorporation were determined by scintillation counting of the fixed samples and the total extractable lipids, respectively. To analyze lipid species, 5,000 counts per minute (CPM) of the lipid sample was spotted at the origin of 100 cm^2^ silica gel 60 aluminum sheets. To separate sulfolipid for quantification, the TLC was developed with a chloroform:methanol:water (90:10:1 v/v/v) solvent system and resolved three times [56]. To separate TAG for quantification, the TLC was developed with a hexane:diethyl ether:acetic acid (80:20:1, v/v/v) solvent system [57]. To examine TDM and TMM accumulation the TLC was developed in a chloroform:methanol:ammonium hydroxide (80:20:2 v/v/v) solvent system. Radiolabeled lipids were detected and quantified using a phosphor screen and a Typhoon Imager, and band density quantified using ImageQuant software [58]. Radiolabeling experiments, lipid extractions and TLCs at pH 5.7 were repeated in at least two independent biological replicates with similar findings in both replicates.

### Measurement of ATP concentration

ATP concentration was measured using the commercially available Cell-Titer Glo kit (Promega) with Relative Luminescence Units (RLU) measured using a Perkin Elmer Envision plate reader. A standard curve of varying ATP concentrations was generated to calculate ATP concentrations of each sample based on RLU measurements, and these concentrations were normalized to bacterial CFU as assayed by viability plating.

### Genetic screens

A transposon mutant library of > 100,000 mutants was generated using the phage Mycomar-T7 as described previously [59]. The library was collected in 4 pools of ∼25,000 mutants, and each pool was plated onto MMAT agar plates (1 g/L KH_2_PO_4_, 2.5 g/L Na_2_PO_4_, 0.5 g/L (NH_4_)_2_SO_4_, 0.15 g/L asparagine, 10 mg/L MgSO_4_, 50 mg/mL ferric ammonium citrate, 0.1 mg/L ZnSO_4_, 0.5 mg CaCl_2_, and 15 g/L agar) containing 10 mM glycerol as a single carbon source and buffered to pH 5.7 with 100 mM MES [55]. Mutants capable of forming colonies on these plates were isolated and confirmed for acidic pH growth in liquid culture (MMAT + 10 mM glycerol buffered to pH 5.7 with 100 mM MES). The transposon insertion sites for confirmed mutants were identified using the inverse PCR technique [60]. In addition to the transposon-based screen, wild type Mtb was also plated in similar conditions and two spontaneous mutants capable of forming colonies were also isolated. Genetic complementation and qPCR studies were performed using methods as described in Zheng *et al.* [34]

### Whole Genome Sequencing

Genomic DNA of selected mutants as well as a wild type control was isolated, DNA libraries constructed, and sequenced using the Illumina MiSeq, in paired end, 250-bp read format (PE250). After the sequencing run, reads were demultiplexed and converted to FASTQ format using the Illumina bcl2 fastq (v1.8.4) script. The reads in the raw data files were then subjected to trimming of low-quality bases and removal of adapter sequences using Trimmomatic (v0.36) [61] with a 4-bp sliding window, cutting when the read quality dropped below 15 or read length was less than 36 bp. The trimmed reads were then aligned to the CDC1551 reference genome using the Burrow-Wheeler Aligner (BWA, [62]). Genome Analysis ToolKit (GATK, [63]) base quality score recalibration, indel realignment, and duplicate removal were applied and SNP and INDEL discovery performed.

### Determination of MBC90 and measurement of antibiotic tolerance

To measure Mtb antibiotic sensitivity in a variety of media, Mtb was seeded in 30 mL of the specified medium in a T75 flask at an initial density of OD_600_ 0.1 and incubated at 37C with 5% CO_2_ for 3 days. After 3 days, cultures were spun down, resuspended in fresh medium of the same kind at OD_600_ 0.1, and 200 μL/well plated in a 96 well plate. 2 μL of antibiotics or SDS was added in 2.5-fold serial dilutions to the plates, and the Mtb incubated at 37°C in 5% CO_2_ for 6 days. 10-fold serial dilutions of the cultures were spot plated on 7H10 + 10% OADC agar plates on day 0 and day 6. The MBC_90_ was determined by the concentration of antibiotic or SDS that produced a one-log change in spot density at day 6 compared to day 0. Using the same experimental design, colony forming units were also measured after treatment with a single concentration of the following drugs: isoniazid (20 μM), rifampin (0.6 μM), and PA-824 (10 μM).

### Statistical methods

All growth curves were performed in biological duplicate and are representative of at least two independent experimental replicates. The error bars for all growth curves represent the standard deviation of a single experiment, although sometimes are too small to see given the consistency of measurement. For experiments measuring Mtb viability, Mtb was cultured in each treatment condition in biological triplicate. The spot plating assays to determine MBC_90_ were performed in biological duplicate and spot plating performed in technical duplicate and are representative data from two separate experiments. For metabolic profiling, five biological replicates were extracted for each treatment, and statistical significance determined using a MANOVA followed by post-hoc pairwise comparisons Bonferroni adjusted for false discovery. Differences were considered significant at p < 0.05. Statistical data are presented in Supplemental Table 1.

## Acknowledgements

We thank Prof. John McKinney for the generous gift of the Δ*icl1/2* mutant and matched wild type control strain used in this study. We are grateful to the MSU RTSF Mass Spectrometry Core Facility staff for assistance in developing methods for extraction and analysis of metabolites. We also acknowledge technical assistance provided by Navanjeet Sahi, Hannah Bodnar, and Emily Juzwiak. JB was supported by a Robert J. Schultz student research award. This study was supported by funding from the Michigan State University start-up funds, AgBioResearch and grants from the NIH-NIAID (U54AI057153 and R01AI116605).

---

**Supplemental Figure 1.**
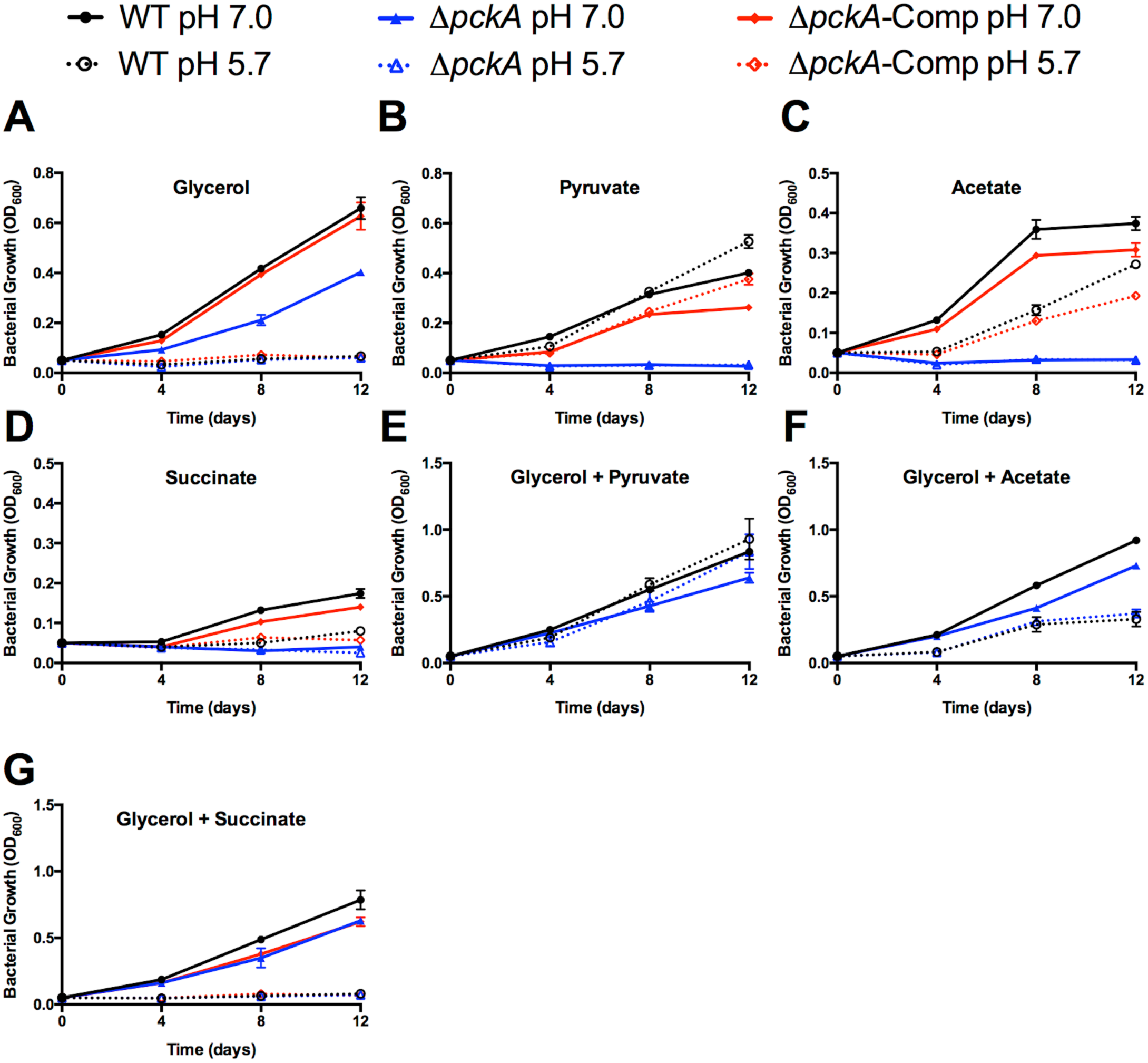
Wild type growth phenotypes are restored in the *ΔpckA* mutant with addition of glycerol. Growth of WT, *ΔpckA* mutant, and complemented strain *(ΔpckA*-Comp) was measured over time in minimal media supplemented with the indicated carbon sources. A-D) *ΔpckA* maintains growth arrest on glycerol at pH 5.7, but is deficient for growth on pyruvate, acetate, and succinate as indicated by a decrease in OD_600_ over time. E-G) Addition of glycerol as a second carbon source restores WT levels of growth in the *ΔpckA* mutant on acetate, pyruvate, and succinate.

**Supplemental Figure 2.**
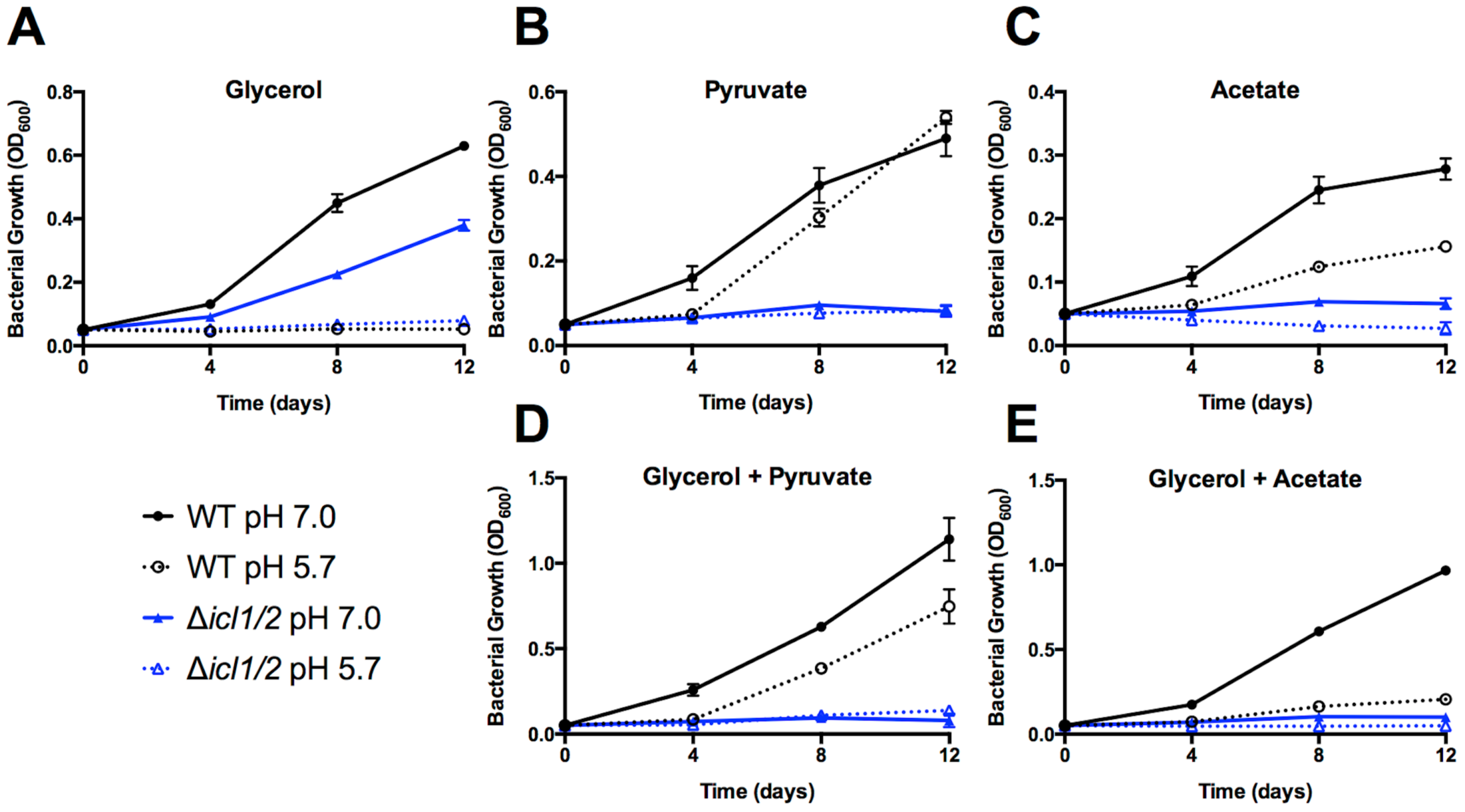
Growth defects of *Δicl1/2* mutant are not restored with the addition of glycerol as a second carbon source. Growth of WT and *Δicl1/2* mutant was measured over time in minimal media supplemented with the indicated carbon sources. A-B) *Δicl1/2* achieves mild bacterial growth on both glycerol at pH 5.7 and pyruvate at pH 7.0 and pH 5.7. This low-level bacterial growth is increased compared to WT at pH 5.7 on glycerol and reduced compared to WT on pyruvate. C) The *Δicl1/2* mutant is unable to grow with acetate as single carbon source. D-E) Addition of glycerol as a second carbon source does not restore growth of the *Δicl1/2* mutant.

**Supplemental Figure 3.**
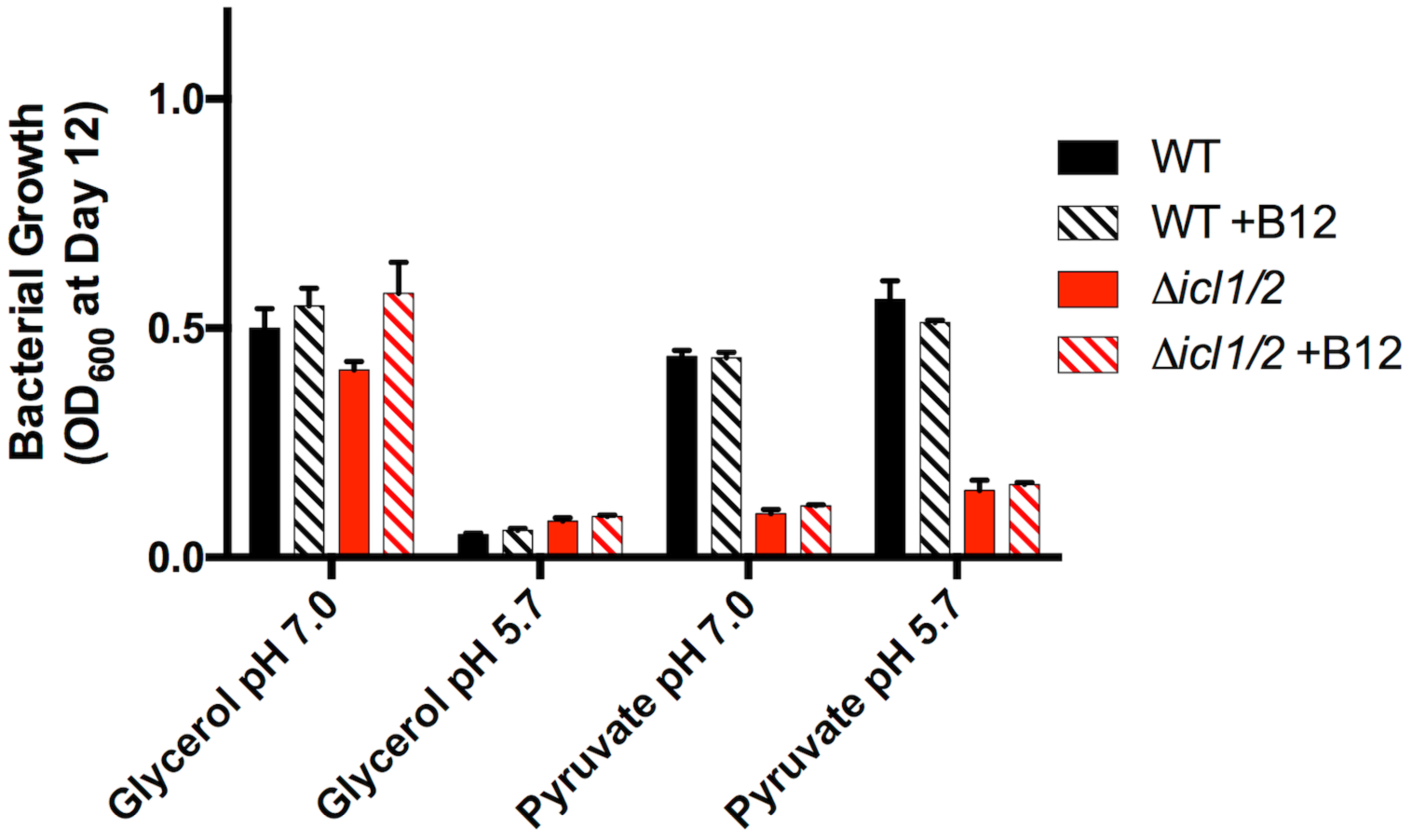
Growth of *Δicl1/2* mutant at acidic pH is not affected by addition of vitamin B12. Summary data showing the OD_6_00 measured on day 12 of growth curves performed in minimal media containing either glycerol or pyruvate as single carbon sources, buffered to pH 7.0 or pH 5.7, and with or without supplementation of vitamin B12. No difference in Mtb growth was observed with supplementation of vitamin B12.

**Supplemental Figure 4.**
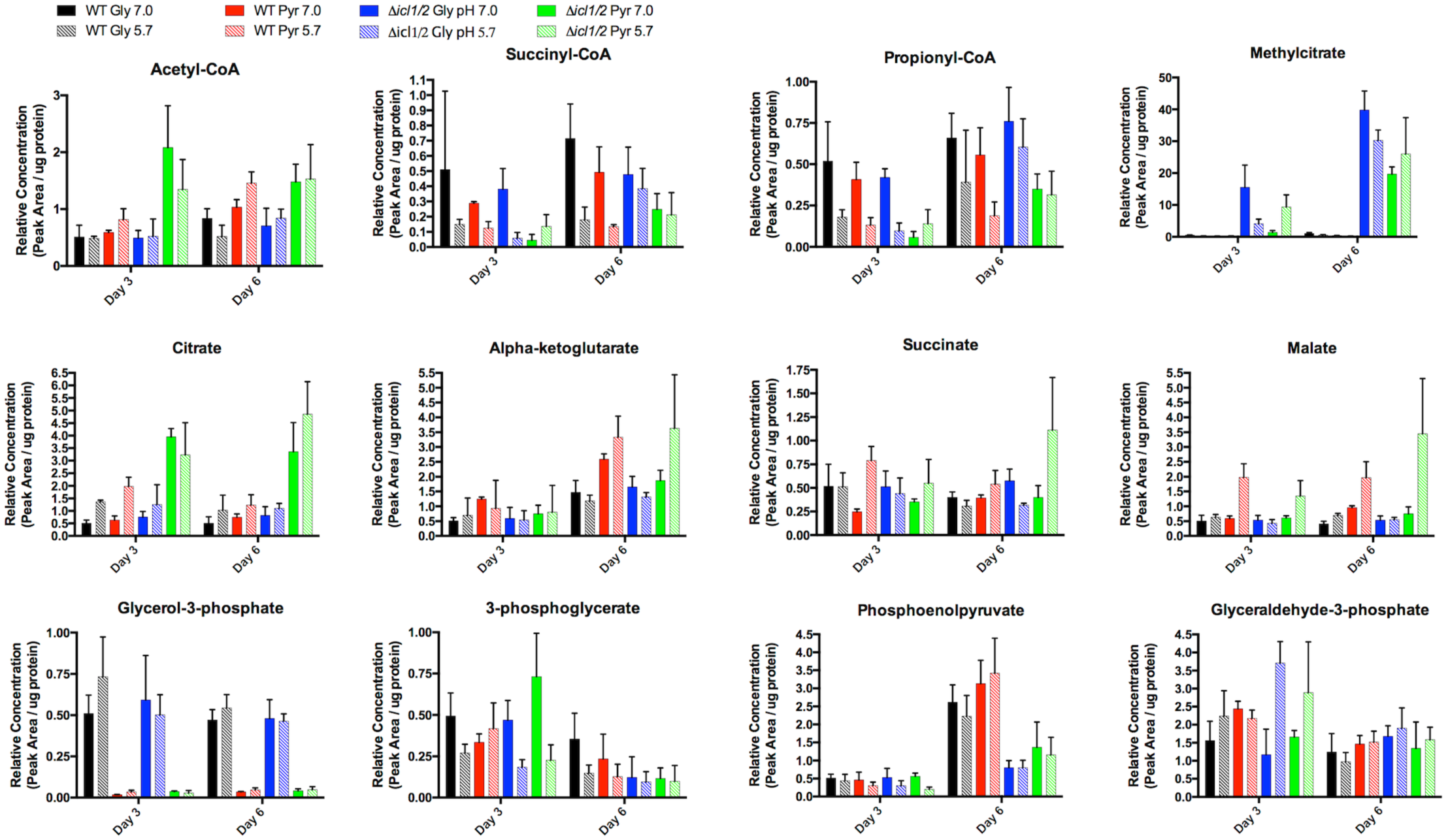
Metabolic profiling of Mtb Erdman wildtype and *Δicl1/2* mutant strains on minimal media agar plates buffered to pH 7.0 or pH 5.7 and containing either glycerol or pyruvate as a single carbon source. Metabolite concentration is reported as the relative peak area per μg of protein for each treatment. Error bars represent the standard deviation. Statistical analyses are included in Supplemental Table 1.

**Supplemental Figure 5.**
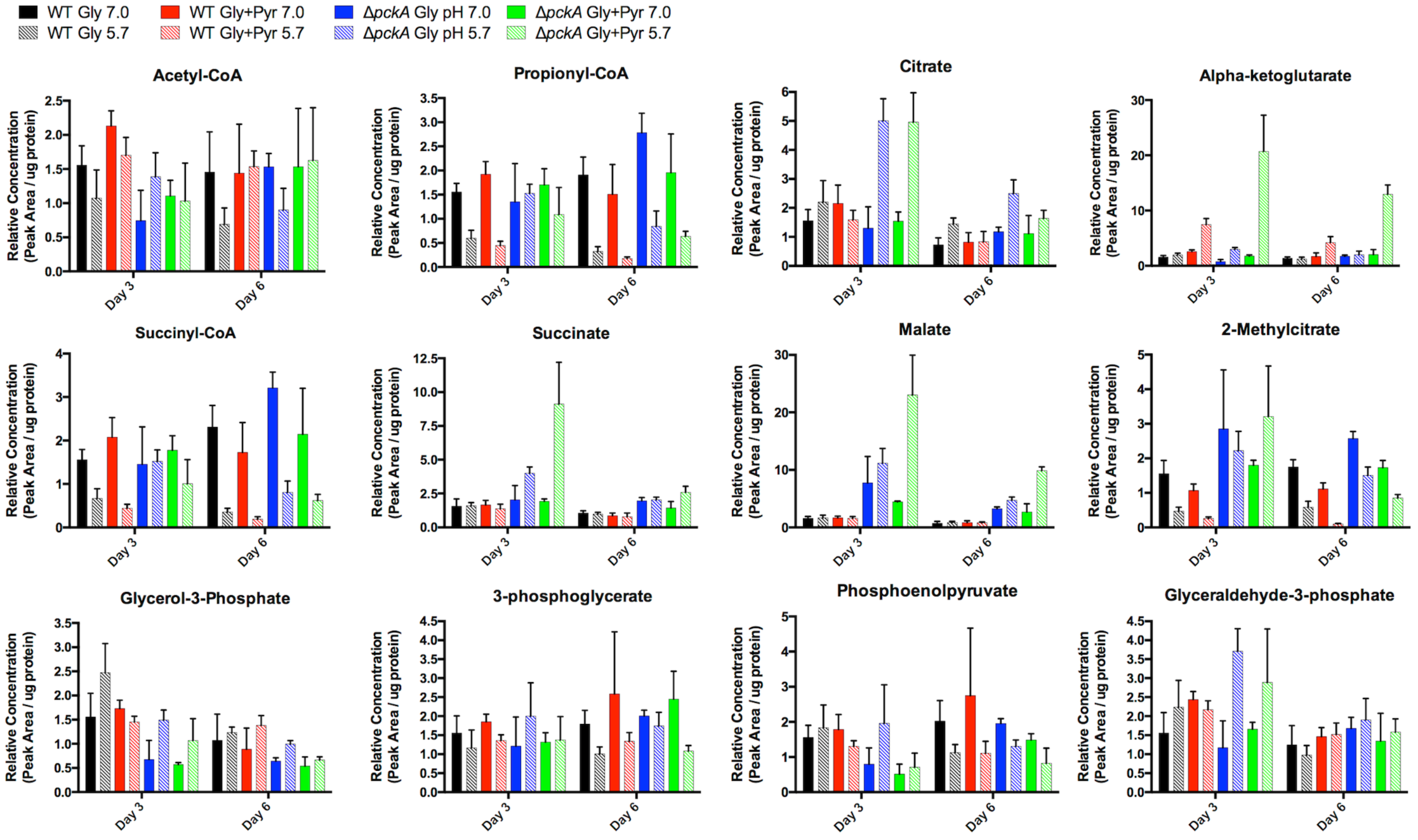
Metabolic profiling of Mtb Erdman wildtype and *ΔpckA* mutant strains on minimal media agar plates buffered to pH 7.0 or pH 5.7 and containing either glycerol or glycerol and pyruvate. Metabolite concentration is reported as the relative peak area per μg of protein for each treatment. Error bars represent the standard deviation. Statistical analyses are included in Supplemental Table 1.

**Supplemental Figure 6.**
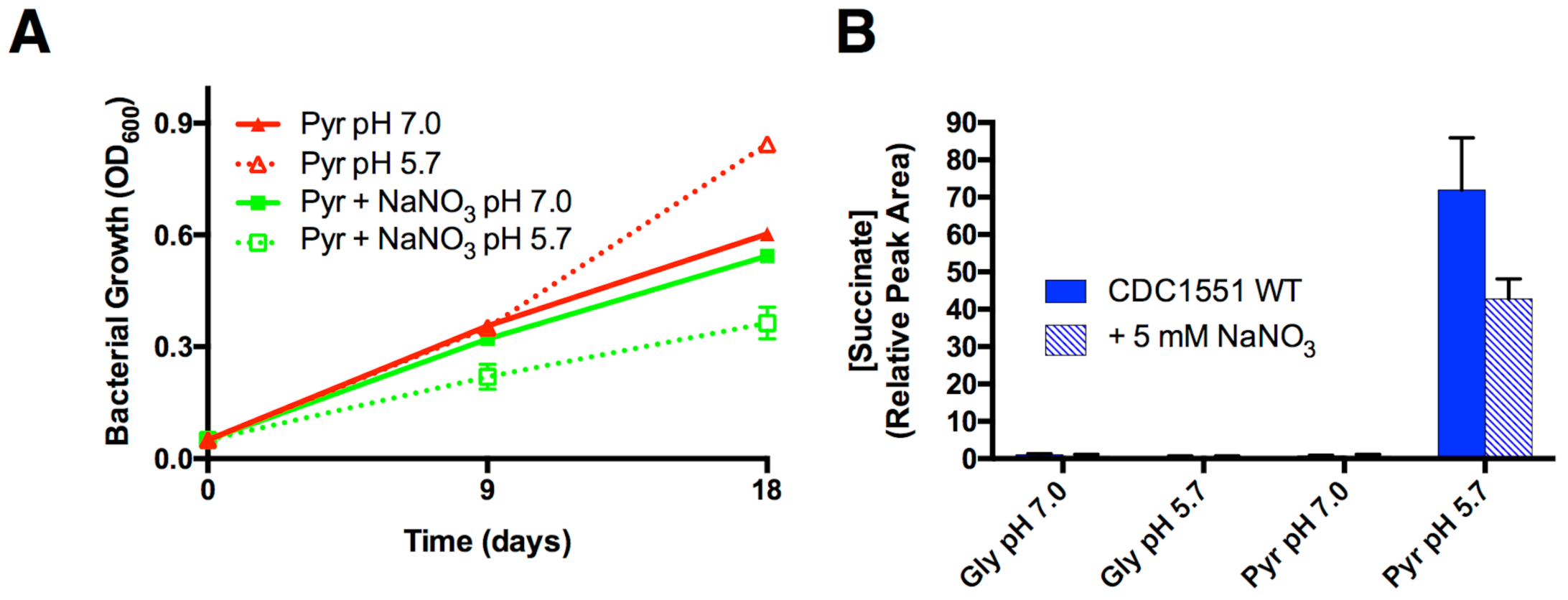
Nitrate decreases Mtb growth on pyruvate specifically at acidic pH but has little effect on succinate secretion. A. Growth of Mtb in minimal medium with pyruvate as a single carbon source buffered to pH 7.0 or pH 5.7 with or without 5 mM sodium nitrate (NaNO_3_). Mtb growth at pH 7.0 is not affected by addition of nitrate, but at pH 5.7 addition of nitrate causes a ∼50% inhibition of growth. **B.** Secretion of succinate is decreased ∼40% with the addition of nitrate at pH 5.7 with pyruvate as a single carbon source, but this could be due to the decreased growth in this condition.

**Supplemental Figure 7.**
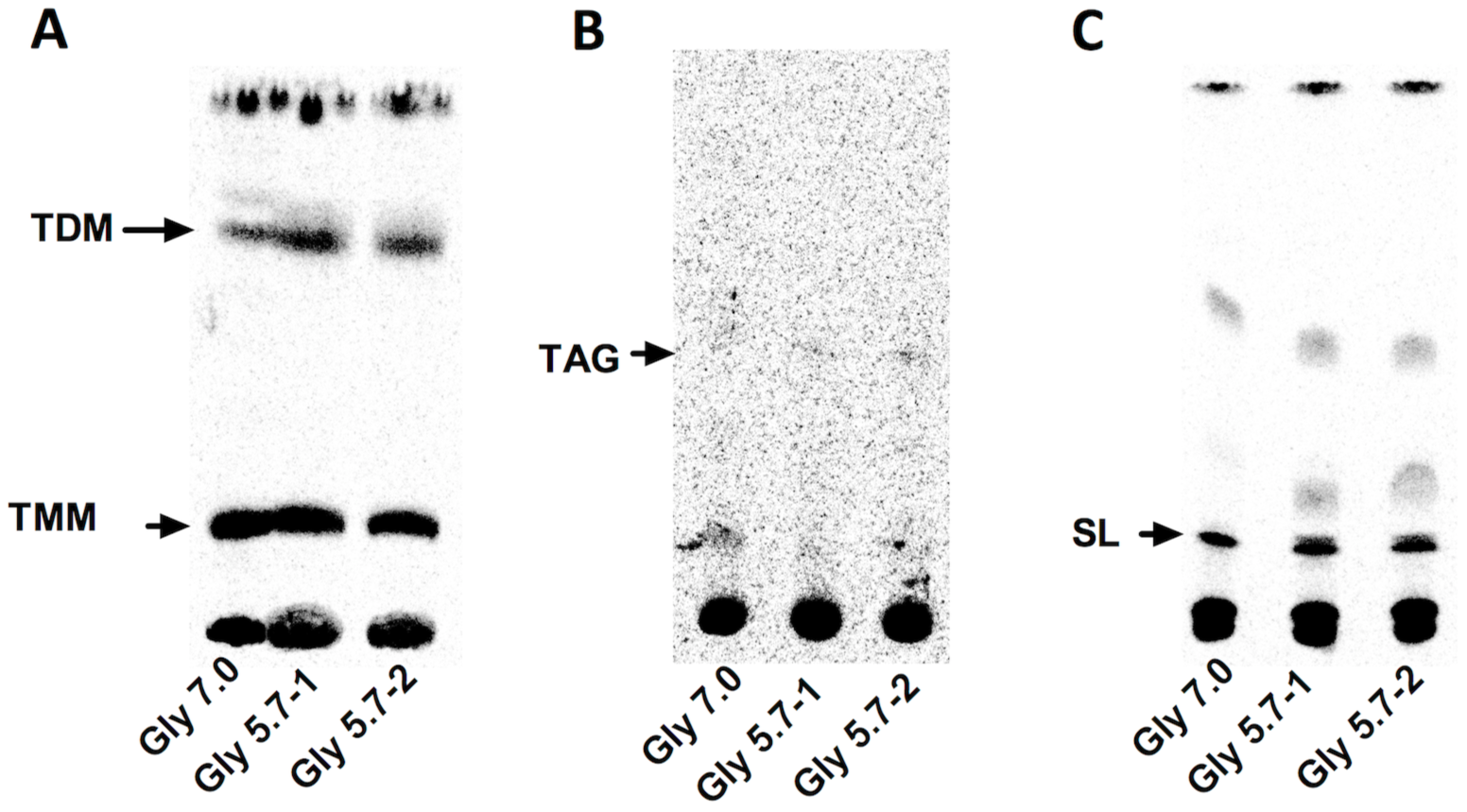
Mtb utilizes glycerol for anabolic metabolism during acid growth arrest. TLC images showing relative abundance of TDM and TMM (**A**), TAG (**B**), and SL (**C**) at pH 7.0 and pH 5.7. Barely detectable quantities of ^14^C-glycerol were incorporated into TAG.

**Supplemental Figure 8.**
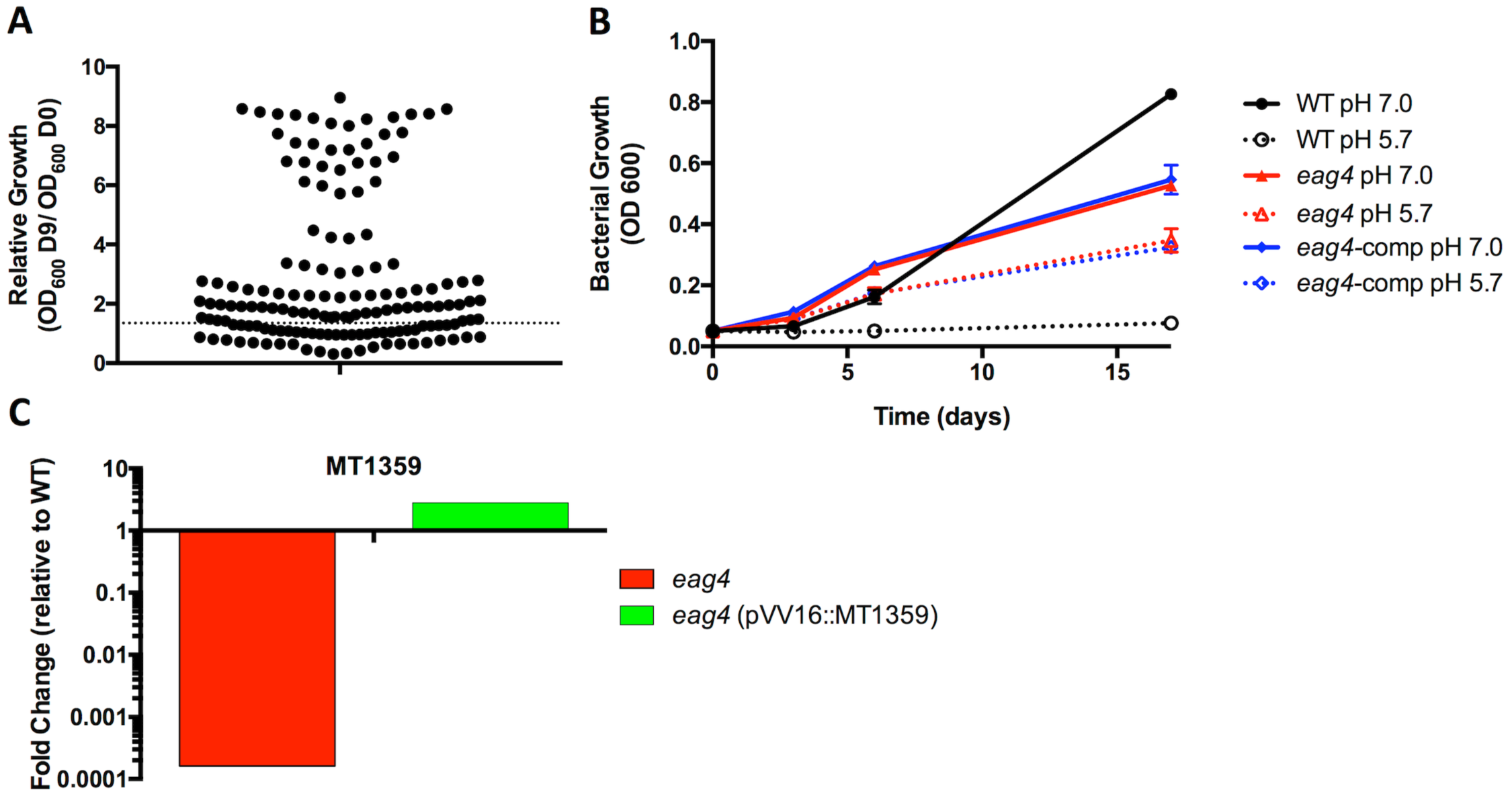
Genetic screen to identify mutants with enhanced growth at acidic pH. Mutants able to form colonies on agar plates buffered to pH 5.7 containing glycerol as a single carbon source were isolated and confirmed as enhanced acid growth (*eag*) mutants by measuring growth in liquid culture conditions of acid growth arrest. **A.** Compiled growth phenotypes for all isolated mutants. Each dot represents an individual mutant, with the fold change in OD_600_ from day 0 to day 9 reported. The dotted line represents the fold change observed in the WT control. **B.** Failed phenotypic complementation of a transposon *eag4* mutant, Tn:MT3159. The mutant was complemented by introduction of an integrative plasmid containing an intact version of the disrupted gene, MT3159, and its native promoter. WT, wild type Mtb. **C.** Genetic complementation of Tn:MT3159. Quantitative real time PCR revealed that the complementation strain (Comp) of Tn:MT3159 can restore the decreased expression of MT3149 transcript levels in the Tn mutant.

